# MODULAR CLICK-CHEMISTRY-BASED ENGINEERING MINIMALLY PERTURBS CRISPR RIBONUCLEOPROTEIN FORMATION FOR GENOME IMAGING APPLICATIONS

**DOI:** 10.64898/2026.09.10.750574

**Authors:** Charlotte Van Tricht, Saphyra Thonon, David Wouters, Alejandro Sifrim, Katy Vandereyken, Vitor B. Pinheiro, Thierry Voet, Jeroen Lammertyn, Dragana Spasic

## Abstract

CRISPR-Cas9 has become a key genome imaging tool, because of its sequence-specific binding to genomic DNA. To achieve signal amplification or multiplexing, various complex CRISPR-Cas9 modifications were introduced, primarily relying on expression-based delivery. However, these engineering strategies often invoke structural perturbations and their performance remained dictated by vector design and in vivo CRISPR ribonucleoprotein (RNP) assembly. To date, research mainly focused on optimizing the expression-based delivery mechanisms of CRISPR components rather than addressing the engineering itself. Here, we developed a modular SPAAC (bio-orthogonal strain-promoted azide-alkyne cycloaddition) click-chemistry-based strategy for engineering the CRISPR RNP that is completely dissociated from its assembly. We demonstrated that CRISPR can be modified (with fluorophores and DNA) at all stages of RNP in vitro assembly and subsequently validated the technology for genome imaging. Importantly, we also revealed that engineering native CRISPR RNP only after it is being formed, minimizes structural perturbations to the complex and yields a two-fold enhanced accessibility of ‘clicked’ DNA. Finally, we utilized SPAAC to conjugate non-native RNA-DNA hybrids, regardless of 5’-3’ directionality, to develop a covalently-attached rolling-circle signal amplification approach. The modularity and universal nature of our approach holds the potential to deliver significant advances to both CRISPR imaging and prime-editing fields.

**GRAPHICAL ABSTRACT:** 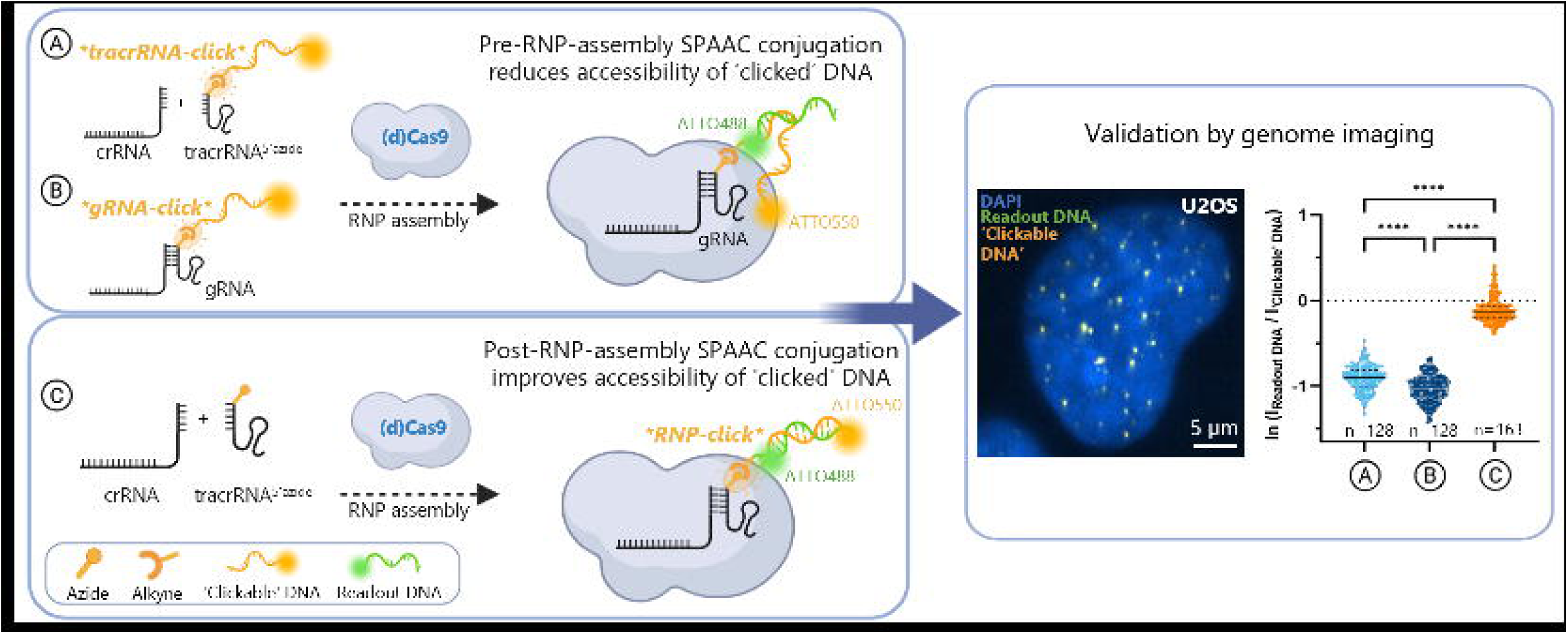

## INTRODUCTION

After more than a decade of research and discovery, various CRISPR-Cas9 (Clustered Regularly Interspaced Short Palindromic Repeats - CRISPR-associated protein 9)-enabled technologies have emerged, ranging from tailored gene-editing strategies (1–5) to biosensing and genome imaging (6, 7). For the latter, CRISPR has been proven a powerful alternative to fluorescence *in situ* hybridization (FISH). Specifically, traditional DNA FISH-based imaging techniques rely on denaturation of genomic double-stranded (ds) to single-stranded (ss) DNA. Moreover, they involve RNAse and protease treatments that do not preserve fragile RNA and protein structures (*i.e.*, chromatin), respectively, thereby complicating its application in multi-omic studies (*e.g.*, genome-transcriptome or genome-proteome studies). In this context, the binding of CRISPR-Cas9 to native genomic DNA and chromatin not only resolves the intrinsic limitations of FISH-based genome imaging but also enables live-cell studies that have been previously unattainable (8).

The CRISPR-Cas9 complex comprises a guide RNA (gRNA) and Cas9 protein that form a ribonucleoprotein (RNP) complex capable of recognizing dsDNA in the presence of a protospacer-adjacent motif (PAM, *e.g*., NGG for *Streptococcus pyogenes* Cas9). Whereas the Cas9 protein cleaves dsDNA, its nuclease-deficient (dead) Cas9 counterpart (i.e., dCas9) only facilitates dsDNA recognition, hence enabling its application in genome imaging studies. The gRNA consists of a CRISPR RNA (crRNA), which specifically recognizes the target through sequence hybridization, and a trans-activating crRNA (tracrRNA) that facilitates recognition of the gRNA by the (d)Cas9 protein.

To unlock the potential of CRISPR-(d)Cas9-guided genome imaging, various fluorescent signal generation approaches have been developed for both fixed- and live-cell imaging. These developments were aimed at achieving improved sensitivity and specificity (often through signal amplification), as well as increased multiplexing (7, 9). As such, increasingly complex and demanding imaging requirements have been achieved through protein and nucleic acid (NA) engineering of the CRISPR-(d)Cas9 RNP. However, those advances have primarily focused on what can be delivered to cells via plasmids and viral vectors. Among those, labelling (d)Cas9 with fluorescent proteins (*e.g.*, enhanced (e)GFP, BFP, mCherry) was foundational for the CRISPR imaging field (10–13). Additionally, one of the first advanced signal amplification approaches, named CRISPR-SunTag (14, 15), relies on (d)Cas9 protein engineering to yield a large ∼1400 kDa structure, offering limited design flexibility and being technically difficult (16). As an alternative, engineering of the gRNA has been adopted widely, owing to the design flexibility, technical simplicity, and commercial availability of engineered NAs (17–20). Most gRNA engineering approaches involve the insertion of protein-recruiting aptamer motifs (*e.g.*, MS2, BoxB, PP7), or larger assemblies thereof, in various locations in the gRNA (17–19, 21–23). However, research suggested that these gRNA-aptamer constructs interfere with crucial internal CRISPR RNP interactions and consequently affect downstream imaging performance (16, 20, 22, 24). Although some non-traditional engineering approaches included the use of protein tags (*e.g.,* Halo, SNAP, BAP), their flexibility remains underexplored, and they failed to gain practical adoption in the genome imaging field (25–27). Moreover, because researchers’ efforts are more focused on optimizing how to deliver CRISPR complex into the cells rather than the CRISPR engineering methodology itself the current CRISPR imaging capabilities remain not fully exploited, often with unaddressed practical challenges (28–30).

In this context, we present an elegant approach for engineering of the tracrRNA of the CRISPR-(d)Cas9 complex through Strain-Promoted Alkyne-Azide Cycloaddition (SPAAC) click-chemistry (**Figure 1**) (31). SPAAC click-chemistry involves a strain-promoted, bio-orthogonal reaction between a constrained alkyne, such as dibenzocyclooctyne (DBCO), and an azide functional group, forming a covalent triazole linkage between two biomolecules without the need for cytotoxic Cu^2+^ catalysts (**Figure S1**) (31). Here, we identify a suitable labelling site in the CRISPR RNP, based on its reported crystal structure. Further, we ‘click’ a fluorophore on the tracrRNA at each of the three CRISPR RNP formation stages (**Figure 1**), followed by validation of both SPAAC click-chemistry and CRISPR functionality using a fluorescent electrophoretic mobility shift assay (fEMSA). Subsequently, we assess the capacity of such engineered CRISPR complexes to bind telomeres in fixed human cells. As ‘click’ partners can be varied, we also investigate whether SPAAC click-chemistry allows CRISPR modification (at each of the three CRISPR RNP formation stages) with a DNA oligonucleotide without interfering with the native CRISPR RNP complex. The accessibility of this ‘clicked’ DNA is then investigated by hybridization of a complementary fluorescent readout probe. Ultimately, as SPAAC click-chemistry allows ‘clicking’ of NAs regardless of their 5’-3’ directionality, we explore how this allows CRISPR to harbor a rolling-circle amplification (RCA) construct, while maximizing the accessibility of the ‘clicked’ RCA primer. Together, we present here an innovative and robust approach for engineering the CRISPR complex with various readout entities and with the additional flexibility to achieve this at its three formation stages. This opens new opportunities for engineering the CRISPR RNP complex in genome imaging as well as genome editing applications.

**Figure 1:**
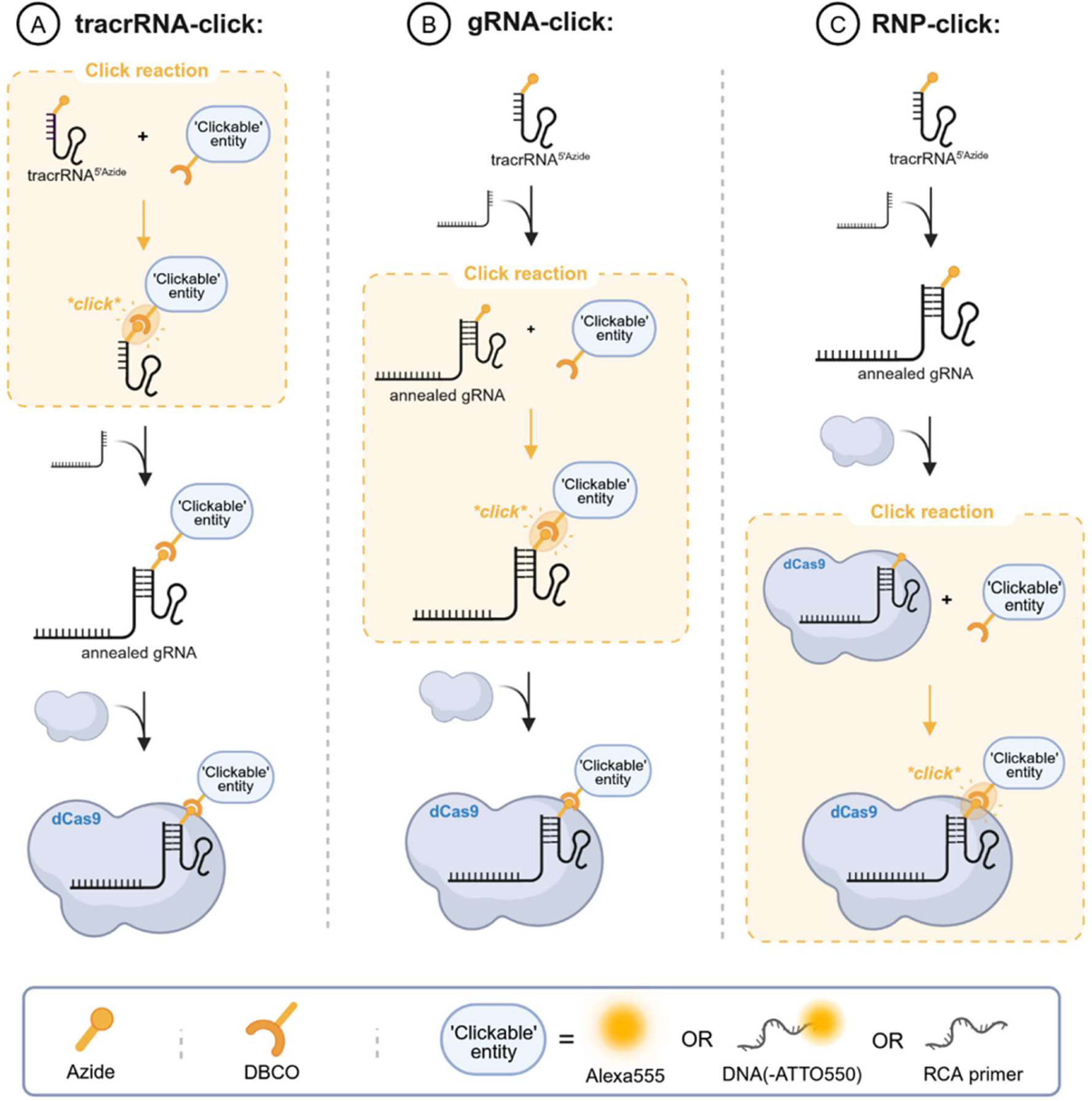
Overview of the engineering methodology employing SPAAC click-chemistry for modifying the CRISPR-(d)Cas9 complex. TracrRNA is modified with an azide group at the 5’ end, which reacts with DBCO-labeled molecules being either Alexa Fluor 555 (i.e., DBCO-Alexa555) or a DNA oligonucleotide (e.g., DNA-ATTO550 or RCA primer). SPAAC click-chemistry engineering of tracrRNA takes place: (A) prior to its annealing with crRNA to form gRNA, followed by gRNA-dCas9 recognition and RNP formation (‘tracrRNA-click’ methodology); (B) at the level of annealed gRNA (‘gRNA-click’ methodology), followed by RNP formation or (C) after RNP complex is fully assembled (‘RNP-click’ methodology). Figure created with BioRender.com.

## MATERIAL AND METHODS

### Materials and reagents

Protein (Cat. #0030108094) and DNA (Cat. #0030108035) Lo-Bind tubes were purchased from Eppendorf (Hamburg, Germany) and 35 mm glass-bottom dishes were obtained from Cellvis (Mountain View, CA, USA; Cat. #D35-14-1.5-N). Amicon® Ultra-15 Centrifugal Filter 100 kDa MWCO was purchased from Millipore (Burlington, MA, USA; Cat. #UFC910024). The human osteosarcoma cell line U2OS was obtained from ATCC (Manassas, VA, USA; Cat. #HTB-96) and DH10β cells were purchased from New England Biolabs (NEB, Ipswich, MA, USA; Cat. #C3019I).

McCoy’s 5A Medium was purchased from Thermo Fisher Scientific (Waltham, MA, USA; Cat. #16600082), HyClone™ Bovine Growth Serum from Cytiva (Marlborough, MA, USA; Cat. #SH30541.03), and LB Miller Broth from VWR (Radnor, PA, USA; Cat. #J106-2KG). Alt-R S.p. dCas9 Protein V3 (Cat. #1081067), Alt-R CRISPR-Cas9 tracrRNA (Cat. #1072532), Alt-R CRISPR-Cas9 tracrRNA-ATTO 550 (Cat. #1075927), IDT Duplex Buffer (Cat. #11-01-03-01), and all custom oligonucleotides (i.e., gBlock dsDNA, crRNA_Telomere_, crRNA_GAL4_, tracrRNA^5’Azide^, DBCO-TEG-DNA-ATTO550, DNA_Hyb_-ATTO488, DBCO-TEG-RCA primer, and DNA_RCA_-ATTO550, **Table S1**) were purchased from Integrated DNA Technologies (IDT, Coralville, IA, USA). DBCO-Alexa555 was bought from Jena Bioscience (Jena, Thuringia, Germany; Cat. #CLK-1290-1). Q5® High-Fidelity 2X Master Mix (Cat. #M0492S), Gibson Assembly Master Mix (Cat. #E2611), SNAP-Surface® 649 (Cat. #S9159S), T4 Polynucleotide Kinase (Cat. #M0201S), T4 DNA Ligase (Cat. #M0202S), Exonuclease III (Cat. #M0206S), Monarch® PCR & DNA Cleanup Kit (Cat. #T1030S), rCutSmart Buffer (Cat. #B6004S), deoxynucleotide triphosphate (dNTP) solution (Cat. #N0447S), and Phi29 DNA polymerase (Cat. #M0269L) were all purchased from New England Biolabs (Ipswich, MA, USA). QIAquick PCR purification kit was from Qiagen (Venlo, Netherlands; Cat. #28106). Yeast tRNA (*S. cerevisiae*) was purchased from Merck (Darmstadt, Germany; Cat. #R8759-100UN). The TelC-Alexa488 PNA-FISH probe was obtained from PNA Bio (Newbury Park, CA, USA; Cat. #F1004). Qubit™ Protein Assay Kit (Cat. #Q33211), 10X RNAse-free Phosphate-buffered saline, pH 7.4 (PBS, Cat. #AM9625), 4′,6-diamidino-2-phenylindole (DAPI, Cat. #D1306), UltraPure™ DNAse-RNAse-Free Distilled Water (hereafter “nuclease-free water” (NF), Cat. #10977015), Ni-NTA resin (Cat. #88221), 6X TriTrack DNA loading dye (Cat. #R1161), Tween-20® (Cat. #J20605.AP), and Pierce^™^ No-Weigh^™^ Dithiothreitol (DTT, Cat. #A39255) were all bought from Thermo Fisher Scientific. Pefabloc (Cat. #11429876001) and Roche blocking reagent (Cat. #10057177103) was purchased from Roche (Basel, Switzerland). Heparin Sepharose 6 Fast Flow was purchased from Cytiva (Cat. #17099825). Dynabeads MyOne Streptavidin C1 (Cat. #65001) and 20X Saline-Sodium Citrate (SSC, Cat. #AM9770) were obtained from Invitrogen (Waltham, MA, USA).

### Cell culture

U2OS cells (ethical approval S64403 and S66605) were cultured in McCoy’s 5A Medium supplemented with 10% HyClone™ Bovine Growth Serum at 37 °C and 5% CO_2_.

### PCR amplification

dsDNA-ATTO488 for the fEMSA was generated by PCR amplification of template gBlock dsDNA, ATTO 488-labeled forward primer, and unlabeled reverse primer (See **Table S1**) using Q5^®^ High-Fidelity 2X Master Mix. 2.5 µL of forward primer (10 µM), 2.5 µL of reverse primer (10 µM), and 1 µL of template dsDNA were added to PCR tube with 25 µL of 2X Master Mix and 19 µL of nuclease-free (NF) water. This mixture was subject to thermocycling: (1) initial denaturation for 30 s at 98 °C, (2) annealing during 30 cycles of 7 s at 98 °C, 20 s at 71 °C, and 10 s at 72 °C, and (3) extension for 2 min at 72 °C. dsDNA was purified using QIAquick PCR purification kit according to the manufacturer’s protocol. **Figure S2A** depicts the dsDNA PCR product before (lane 1) and after purification (lane 2).

### CRISPR-dCas9 fEMSA

The CRISPR-dCas9 fEMSA protocol was adapted from Liu and colleagues (32, 33). The dsDNA-ATTO488 concentration required for the assay was determined by titration (**Figure S2B**). gRNA was annealed by incubating 5 µM crRNA and tracrRNA in IDT Duplex Buffer at 95 °C for 1.5 min, followed by gradual cooling to room temperature (RT) over 45 min. Annealed gRNA (1 µL, 5 µM), dCas9 protein (1 µL, 5 µM), and CRISPR reaction buffer (1 µL, 10X: 1 M NaCl, 100 mM MgCl_2_, 500 mM Tris-HCl, pH 7.4) were combined with 7 µL NF water and incubated for 15 min at RT to form the RNP complex, followed by addition of 3 µL purified dsDNA-ATTO488 target and a further 30 min incubation at RT. Samples were mixed with 6X TriTrack DNA loading dye and resolved by agarose gel electrophoresis (120 V, 60 min). Gels were imaged on a Typhoon FLA 9500 scanner (GE Cytiva) using filter sets BPB1 (530DF20) and BPG1 (570DF20) (Fujifilm).

### SPAAC click-chemistry

SPAAC click-chemistry reactions were carried out in an RNAse-free environment and were adapted from Taemaitree and colleagues to accommodate the work presented in this publication (34). For the ‘tracrRNA-click’ method, 3.6 µL of tracrRNA-5’Azide (100 µM) was added to 2.4 µL of HEPES (100 mM), 0.2 µL MgCl_2_ (500 mM) and 1.2 µL KCl (1 M) in a protein Lo-Bind tube. Subsequently, 1.25 µL DBCO-modified molecules (*i.e.,* DBCO-Alexa555, DBCO-TEG-DNA-ATTO550, or DBCO-TEG-RCA primer; 1 mM) was added and incubated at RT for 4 h. The solution was stored at −20 °C until further use.

For the ‘gRNA-click’ method, 3.6 µL crRNA and 3.6 µL tracrRNA-5’Azide were added to 2.4 µL of HEPES (100 mM), 0.2 µL MgCl_2_ (500 mM) and 1.2 µL KCl (1 M), and annealed at 95 °C for 1.5 min, followed by gradual cooldown to RT for 45 min. Subsequently, 1.25 µL DBCO-(TEG)-modified molecule (1 mM) was added and the solution incubated at RT for 4 h, followed by storage in −20 °C until further use.

For the ‘RNP-click’ method, gRNA annealing was first performed as explained in the previous paragraph. Subsequently, 5.8 µL of Alt-R *S.p.* (d)Cas9 protein (62 µM) was added and incubated for 15 min at RT to form a functional CRISPR RNP complex. The SPAAC click-chemistry reaction was initiated by adding 1.25 µL of DBCO-modified molecule (1 mM) followed by incubation of 4 h at RT, after which the molecules were stored at −20 °C until further use.

### CRISPR-based genome imaging in fixed cells

Cells were seeded in 35 mm glass-bottom dishes at a concentration of 37,500-50,000 cells per 2 mL. At 70% confluency, the cells were fixed for 20 min at −20 °C in ice-cold methanol-acetic-acid solution (MAA) at 1:1 ratio. Fixation was followed by three washing steps of 3-5 min in PBS (1X) solution and subsequent storage at 4 °C in PBS (1X). For each experiment, three such dishes served as three technical replicates.

The fixed cells were incubated with pre-assembled RNP following the CASFISH protocol (25). First, the dishes with fixed cells were pre-heated to 37 °C, followed by adding 400 µL of CRISPR binding-blocking buffer (*i.e.*, 20 mM HEPES pH 7.5, 100 mM KCl, 5 mM MgCl_2_, 5% glycerol, 1% BSA, 0.1% Tween-20, and freshly added 1 mM DTT and 0.1 mg/mL yeast tRNA) for 15 min at 37 °C. Then, 400 µL of 5 nM CRISPR complex was added to the cells and incubated for 1 h at 37 °C. CRISPR incubation was followed by three washing steps of 5 min, each using CRISPR binding-blocking buffer. Ultimately, genomic DNA was stained with 6 µM DAPI in PBS for 10 min, followed by three rinses with PBS (1X). Imaging was performed in PBS (1X).

For hybridization of fluorescent probe DNA_Hyb_-ATTO488 to DBCO-DNA-ATT550-labeled CRISPR complexes in situ, DAPI staining was followed by 30 min incubation of 5 nM DNA_Hyb_-ATTO488 in PBS (1X) at RT. After incubation, cells were washed three times with PBS (1X) at RT, and imaging was performed in PBS (1X).

### Combined CRISPR and peptide nucleic acid (PNA)-FISH

For combined CRISPR and PNA-FISH imaging, the former was executed according to the protocol described in the previous section until the washing steps with binding-blocking buffer. Following these washing steps, the cells were kept at 37 °C. Subsequently, the cells were inubated with 50 nM of telomere TelC-Alexa 488 PNA-FISH probe in PNA-FISH hybridization buffer (*i.e.*, 20 mM Tris pH 7, 20% formamide, 0.5% Roche Blocking reagent) at 37 °C for 10 min, followed by additional incubation of 2 h at RT in the dark. This hybridization was followed by a 10 min washing step using the wash solution (2X SSC, 0.1% Tween-20) at RT. Finally, nuclei were stained with DAPI (6 µM) for 10 min, followed by three consecutive washes (2 min each) with SSC (2X), SSC (1X) and NF water. Cells were stored at 4 °C until further use and imaging was performed in PBS (1X).

### Cas9-SNAP cloning

To obtain Cas9-SNAP, the SNAP gene was first introduced into the pLentiCRISPR-Cas9 construct(35), the latter being a gift from Prof. Dirk Daelemans (KU Leuven). The SNAP gene was amplified from the LppOmpA-SNAP construct (36) using primers AA_CS_021 and AA_CS_022 (**Table S1**). The amplified SNAP gene was subsequently inserted into the pLentiCRISPR-Cas9 construct using the Gibson Assembly Master Mix according to the manufacturer’s protocol, resulting in construct pLentiCRISPR-Cas9_Cterminal-SNAP containing the gene of the Cas9-SNAP fused protein.

Next, for expression and purification of the Cas9-SNAP protein, the Cas9-SNAP gene was cloned into the pNGAL97 vector (37). The Cas9-SNAP gene was amplified from pLentiCRISPR-Cas9_Cterminal-SNAP using primers AA_CS_035 and AA_CS_036. The amplified Cas9-SNAP gene was then inserted into the pNGAL97 vector using the Gibson Assembly Master Mix according to the manufacturer’s protocol, resulting in construct pNGAL_Cas9_Cterminal-SNAP. Finally, the construct was transformed into DH10β cells for Cas9-SNAP production.

### Cas9-SNAP expression, purification, and labelling

In order to produce Cas9-SNAP(-Dy649), 2 liter (L) of LB Miller Broth (25 g/L) supplemented with 100 mg/L ampicillin was inoculated with 2.5% of an overnight culture of Cas9-SNAP-expressing DH10β cells, and grown in a 5 L baffled bottom shake flask at 37 °C at 120 rpm. Once the optical density (OD600) reached 0.6, Cas9-SNAP expression was induced with 100 ng/mL anhydrotetracycline for 20 h at 20 °C and at 120 rpm. Cells were harvested by centrifugation (4,000 × g for 10 min at 4 °C) and the pellet was stored at −20 °C.

Subsequently, stored cell pellets were thawed on ice and resuspended in cold heparin binding buffer (100 mM Tris-HCl (pH 8), 20 mM NaCl, 4 mM DTT, 5% (v/v) glycerol, 1 mM Pefabloc). The cells were lysed through sonication (12 rounds of 10 s sonication with 20 s interval) on ice and the soluble fraction was collected after centrifugation (4,000 × g for 15 min at 4 °C). Heparin Sepharose 6 Fast Flow resin slurry (10 mL) was pretreated with 50 mL NaOH (0.1 M) wash solution and equilibrated with 50 mL cold heparin binding buffer. Then, the soluble fraction of the cell lysate was mixed with the Heparin slurry and incubated for 30 min at RT while shaking. The mixture was loaded onto a gravity column and buffer was discarded. The column was washed three times with 50 mL cold heparin wash buffer (100 mM Tris-HCl (pH 8), 100 mM NaCl, 4 mM DTT, 5% (v/v) glycerol) and proteins were eluted with 30 mL of cold heparin elution buffer (100 mM Tris-HCl pH (8), 500 mM NaCl, 2 mM DTT, 5% (v/v) glycerol, 1 mM Pefabloc). For the second purification step, heparin eluate was mixed with 8 mL of Ni-NTA resin slurry (equilibrated with 50 mL of 50 mM Tris-HCl (pH 8) for 30 min at RT) and 22 mL of cold 50 mM Tris-HCl (pH 8), and allowed to bind for 30 min at RT. The mixture was loaded onto a gravity column and buffer was discarded. The column was washed three times with 50 mL cold Ni-NTA wash buffer (50 mM Tris-HCl (pH 8), 100 mM NaCl, 1 mM DTT, 10 mM imidazole (Acros), 5% (v/v) glycerol) and proteins were eluted with 30 mL cold Ni-NTA elution buffer (50 mM Tris-HCl (pH 8), 100 mM NaCl, 1 mM DTT, 500 mM imidazole, 5% (v/v) glycerol, 1 mM Pefabloc). Purified proteins were concentrated and buffer was exchanged to 2X Cas9 Storage buffer (20 mM Tris-HCl (pH 7.4), 600 mM NaCl, 0.2 mM EDTA, 1 mM DTT, 2 mM Pefabloc) using Amicon® Ultra-15 Centrifugal Filter 100 kDa MWCO. Proteins were stored in 1X Cas9 Storage buffer supplemented with 50% glycerol.

Cas9-SNAP was labeled with the SNAP substrate by adding a 1:3 molar excess of 250 μM SNAP-Surface® 649 in DMSO to the protein solution. The solution was then incubated in the dark for 30 min at 37 °C while shaking at 300 rpm. Labeled proteins were purified from unlabeled substrate with Amicon® Ultra-15 Centrifugal Filter 100 kDa MWCO. Cas9-SNAP-649 proteins were stored in 1X Cas9 Storage buffer supplemented with 50% glycerol at −20 °C. Protein purity was verified using 12% SDS-PAGE (**Figure S4**). Concentration of pre-labeled Cas9-SNAP proteins was measured using the Qubit Protein Assay Kit on the Qubit fluorometer. Cas9-SNAP-649 concentrations were estimated by measuring the absorbance of the SNAP-Surface® 649 substrate at 655 nm.

### Dumbbell RCA template design and synthesis

The dumbbell RCA template design (**Table S1**) was modified from an earlier described template (38) by adding 5 DNA bases to the dumbbell stem structure, allowing for the division of the template into two separate stem-loop dumbbell building blocks: one containing the RCA primer binding site (i.e., BB_PrimerBinding_, **Table S1**) and one encoding the fluorescent readout probe binding site (i.e., BB_Readout_, **Table S1**) or a negative control version with nonsense binding site (i.e., BB_Nonsense_). The dumbbell building blocks anneal to each other in a pre-defined dumbbell shape (see **Figure S8A**).

For synthesis, the dumbbell building blocks (*i.e.*, BB_PrimerBinding_ and BB_Readout_ for most experiments, or BB_PrimerBinding_ with BB_Nonsense_ for negative control reactions) were first phosphorylated by mixing 2 μM of each dumbbell building block, T4 DNA Ligase Reaction Buffer (1X) and 10 units of T4 Polynucleotide Kinase in 50 μL reaction volumes. Reactions were incubated for 30 min at 37 °C followed by heat inactivation for 20 min at 65 °C. Dumbbell building blocks were annealed by heating the reactions to 95 °C for 5 min and then gradually cooled to RT. After cooling, 1 μL of T4 DNA Ligase was directly added to the reaction and incubated for 30 min at 37 °C, and the reaction was purified using the Oligonucleotide Cleanup Protocol of the Monarch® PCR & DNA Cleanup Kit. Finally, fully formed dumbbell RCA templates were isolated by adding 20 units of Exonuclease I and 100 units of Exonuclease III to the reaction supplemented with rCutSmart Buffer (1X). Reactions were incubated for 30 min at 37 °C and purified with the Oligonucleotide Cleanup Protocol of the Monarch® PCR & DNA Cleanup Kit.

### Standard in vitro RCA reactions

1 μL of RCA primer (10 μM, **Table S1**), 2 pmol of dumbbell RCA template, 0.5 μL dNTP mix (10 mM) and Phi29 DNA Polymerase Reaction Buffer (1X) were mixed in a total reaction volume of 13 μL. The RCA primer was annealed to the dumbbell RCA template (heating to 95 °C for 5 min and then gradually cooled to RT) followed by the addition of 1.5 μL Recombinant Albumin and 0.5 μL Phi29 Polymerase. RCA reactions were incubated for 3 h at 30 °C and heat inactivated for 20 min at 60 °C. **Figure S8B** shows verification of RCA products and corresponding negative control conditions.

### In vitro RCA with fluorescent readout probes

5 μL of Dynabeads were washed three times and blocked for 30 min at RT with 500 μL of binding-washing buffer (5 mM Tris-HCl (pH 7.5), 0.5 mM EDTA, 1 M NaCl, 0.01% Tween-20). RCA products were captured by incubation with the blocked Dynabeads in 100 μL of binding-washing buffer for 30 min at RT without shaking or rotation. Dynabeads were carefully washed (without vortexing) three times in 0.01% Tween-20. Template and non-biotinylated products were washed away by adding 500 μL of NaOH (10 mM) for 30 s, followed by a quick wash with 500 μL of NaOH (10 mM) and three washes with 500 μL of 0.01% Tween-20.

For probe annealing, beads were resuspended in 16 μL of binding-washing buffer and RCA product was labeled by annealing 4 μL of RCA fluorescently labeled probes (10 μM) at 75 °C for 5 min before gradually cooling to RT. Beads were carefully washed three times with 500 μL of 0.01% Tween-20 and labeled RCA products were eluted with 20 μL of 1 mM biotin elution buffer (10 min incubation at RT followed by 5 min incubation at 99 °C). End-point fluorescence intensities (**Figure S8C**) were measured by using the CLARIOstar microplate reader (BMG Labtech) with excitation at 483 nm and emission at 530 nm. 1 mM biotin elution buffer was used as a blank measurement.

### In situ CRISPR-RCA with fluorescent readout probes

The standard CRISPR-based imaging protocol as described under *CRISPR-based genome imaging in fixed cells* was further adapted to accommodate the RCA reaction. CRISPR complex incubation and final binding-blocking buffer washes were followed by dumbbell RCA template incubation (0.4 µL of 42.4 ng/µL template in 100 µL 1X Phi29 buffer) at 37 °C for 45 min. Excess dumbbell RCA template was removed by three rinses with 200 µL of CRISPR binding-blocking buffer. Subsequently, RCA solution (1 µL dNTPs (1 mM), 0.6 µL Phi29 polymerase, and 1 µL BSA in 97 µL of Phi 29 buffer (1X)) was added for 2 h at 30 °C to facilitate the RCA reaction, followed by three washes with CRISPR binding-blocking buffer (supplemented with 6 µM of DAPI) for 5 min each. Finally, fluorescent readout probe (25 nM in PBS (1X)) was incubated for 30 min at 37 °C, followed by three washes with PBS (1X) and fluorescence readout in PBS (1X).

### Image acquisition and pre-processing

Fluorescence imaging of fixed cells was performed using widefield Nikon TI Eclipse microscope (Nikon Instruments). Z-stacks were acquired with a stepsize of 0.2 μm, covering a total thickness of about 8 μm, which is enough to image the entire nucleus of the cell.

Afterwards, fluorescence images were subject to linear image processing using Nikon NIS-Elements (version 6.20.02) General analysis 3 software. Concretely, out-of-focus fluorescence signal was removed by the NIS-Elements ‘Clarify.ai’ tool. Then, 3D image stacks were processed by the ‘Extended Depth of Focus’ tool to compile the 3D stack into a 2D projection, and sharpness improved with ‘Gauss-Laplacian’ tool (default value). The images included in this work were all subject to this same image processing workflow. Subsequently, fluorescent foci were identified automatically by running the acquired images through nuclei segmentation and fluorescent foci detection protocols (see next section).

### Nucleus segmentation for fluorescent foci detection

For visual guidance on image processing, please refer to **Figure S10**. The pre-processed ‘.nd2’ images were first split into separate channels using nd2reader (version 3.3.0). Individual nuclei were segmented with Cellpose (version 2.1.0) using the ‘nucleus’ model and a diameter parameter of 200 (39). Nuclei located on the border of the acquired field of view were eliminated as they do not represent a complete nucleus, and thus would distort the results. Nuclei consisting of less than 5000 pixels were also left out, since they were unusually small and were most likely segmentation errors. The segmentation boundary was expanded by 10 pixels (∼0.65 μm) using Scikit-image (version 0.19.3., skimage.segmentation.expand_boundaries), as Cellpose tended to slightly underestimate the size of nuclei (40). Single nuclei images for each imaging channel were then made by extracting the bounding box for each nucleus, along with a mask of the segmentation boundary, which was used to filter out fluorescent foci unassigned to nuclei. The channels containing the fluorescent foci were then subjected to a Gaussian filter (skimage.filters.gaussian, sigma=1) to remove any noise induced by the microscope.

Fluorescent foci detection was done on a nucleus-by-nucleus basis using scikit-image’s implementation of the Laplacian of Gaussian algorithm (skimage.feature.blob_log), the performance of which is determined by 3 parameters: min_sigma and max_sigma, that effectively define how small/large fluorescent foci are allowed to be, and threshold which determines how bright fluorescent foci have to be for it to be considered as such. Parameters were adjusted between the processing of each separate dataset (see Table S2 for the full overview), but generally were set as: min_sigma=2, max_sigma=3, threshold=0.001. In case multiple fluorescence imaging channels were present, colocalization of fluorescent foci between channels was calculated by matching them using bigfish.multistack.detect_spots_colocalization with a threshold of 10.

Prior to plotting and statistical analysis of the extracted fluorescent foci data, each nucleus was subject to visual validation to include missed (or omit incorrect) fluorescent foci. Concretely, we eliminated (1) aberrant nuclei (*e.g*., with highly unusual shapes), (2) nuclei from which the 3D volume was not entirely captured by the acquired Z-stack (*i.e*., incomplete data), and (3) nuclei which exhibited clear signs of mitosis (*e.g*., metaphase chromosomes).

### Statistical analysis

GraphPad Prism 10 (version 10.2.3, GraphPad Software) software was utilized for statistical analysis and data visualization. Statistical analysis of fluorescent foci counts was performed using non-parametric Kruskal-Wallis test followed by Dunn’s multiple comparisons post-hoc test, as the count data was discrete, variances were statistically different, and datasets were reasonably large. For the hybridization efficiency ratio in Figure 5D, logarithmic transformation was performed on the ratio data to account for inherent skewness, and to satisfy variance assumptions for parametric tests. Ordinary one-way ANOVA with Tukey’s multiple comparisons post-hoc test was performed on this natural log-transformed dataset to assess statistical significance among the groups. Experiments were carried out with three technical repetitions and data were pooled for final analysis. All statistical tests were two-tailed, with a significance threshold set α< 0.05.

## RESULTS

### SPAAC click-chemistry enables modification with a fluorophore at three stages of CRISPR RNP formation

To engineer the CRISPR-(d)Cas9 complex using SPAAC click-chemistry, we relied on its reported crystal structure that reveals a protrusion of the crRNA-tracrRNA duplex outside of the RNP complex (**Figure 2A**) (41). We hypothesized that modifying the tracrRNA with an azide functional group at its 5’-end would lead to a spatially accessible site for labelling by SPAAC click-chemistry. Here, tracrRNA was selected as the invariable 67-mer component of the CRISPR-(d)Cas9 complex, as opposed to target-specific crRNA, with the aim to simplify the approach. To test our hypothesis, we ‘clicked’ a DBCO-carrying molecule (so-called “clickable entity”) at three different stages of CRISPR RNP complex formation (**Figure 1**), namely at the stage of: (a) the tracrRNA, prior to gRNA annealing and subsequent full RNP formation (‘tracrRNA-click’), (b) the annealed gRNA, followed by assembly with the dCas9 protein into a full RNP complex (‘gRNA-click’) or (c) the assembled CRISPR RNP (‘RNP-click’).

**Figure 2.**
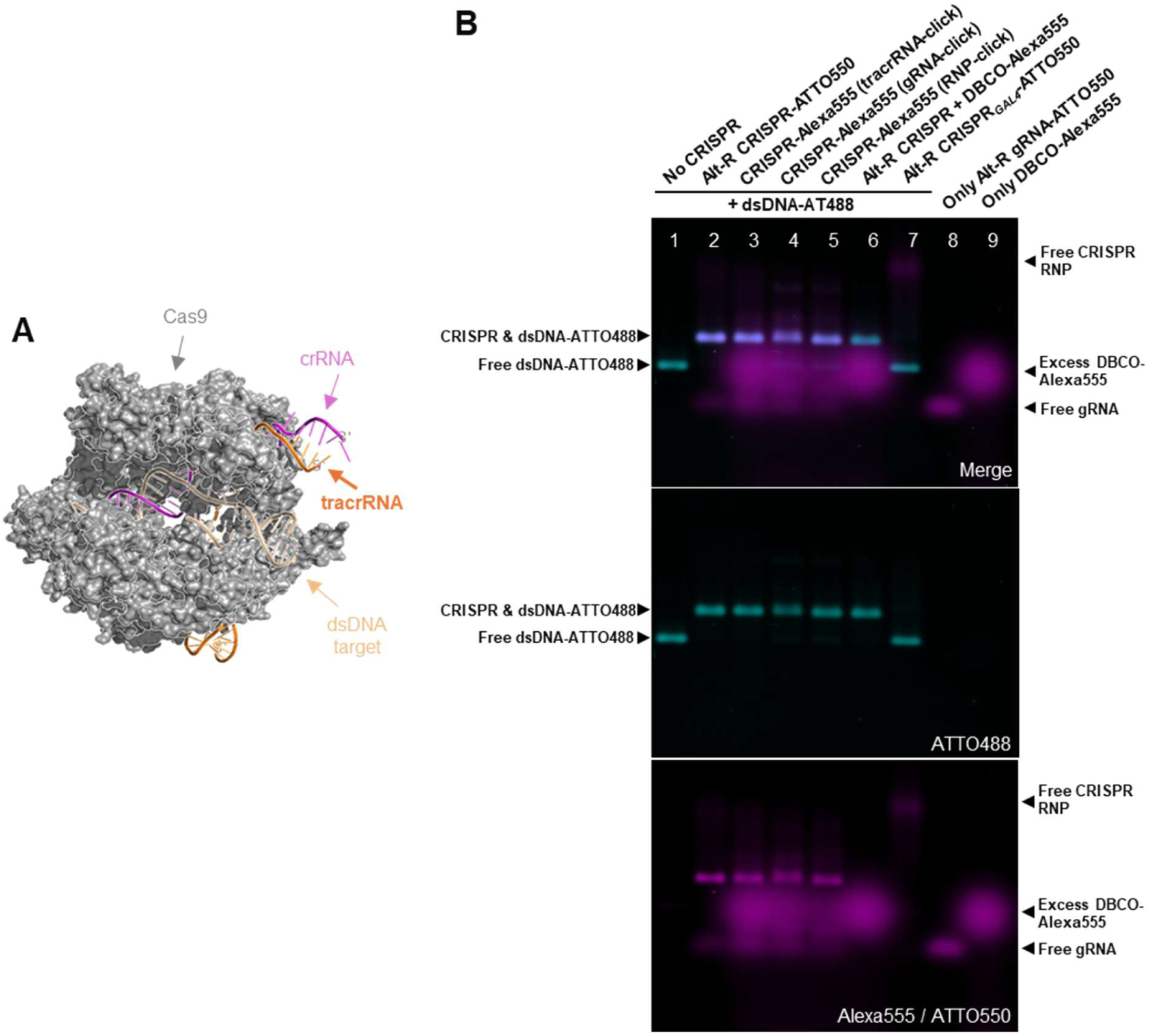
SPAAC click-chemistry enables CRISPR engineering with a fluorophore at all three stages of RNP formation. **(A)** 3D render of the CRISPR-Cas9 bound to an arbitrary dsDNA target (Protein Databank, 8KAH). The tracrRNA-crRNA duplex protrudes out of the RNP structure. 3D render created with PyMOL (pymol.org). **(B)** fEMSA demonstrates fluorophore SPAAC click-chemistry and CRISPR binding to a fluorescent 433 bp dsDNA-ATTO 488 (dsDNA-ATTO488) with telomere-like target sequence. The ‘Merge’ panel shows (colocalized) fluorescence of dsDNA-ATTO488 (cyan, ‘ATTO488’ panel) and Alexa Fluor 555 or ATTO 550 (magenta, ‘Alexa555 / ATTO550’ panel) SPAAC-‘clicked’ CRISPR complexes, as well as positive and negative control CRISPR complexes, binding to the dsDNA-ATTO488 target. Lane 1: reference dsDNA-ATTO488 in absence of CRISPR complex (‘No CRISPR’). Lane 2: positive control showing a native CRISPR complex, with fluorescently labeled Alt-R tracrRNA-ATTO550, targeting the telomere-like dsDNA-ATTO488 (‘Alt-R CRISPR-ATTO550’). Lanes 3-5: the SPAAC click-chemistry reaction with DBCO-Alexa555 according to the three engineering methodologies (i.e., tracrRNA-click, gRNA-click, and RNP-click) as depicted in Figure 1 (‘CRISPR-Alexa555’). Lane 6: a native CRISPR complex with Alt-R tracrRNA, in absence of azide modification and in presence of DBCO-Alexa555, binding to dsDNA-ATTO488 (‘Alt-R CRISPR + DBCO-Alexa555’). Lane 7: GAL4-targeted CRISPR complex with Alt-R tracrRNA-ATTO550 that is unable to bind dsDNA-ATTO488 (‘Alt-R CRISPR_GAL4_-ATTO550’). Lanes 8-9: free fluorescent gRNA with Alt-R tracrRNA-ATTO550 (‘Only Alt-R gRNA-ATTO550’) and excess DBCO-Alexa555 (‘Only DBCO-Alexa555’), respectively.

To assess effectiveness of these three SPAAC click-chemistry approaches and the subsequent ability of CRISPR RNP to form a functional ternary complex with target dsDNA, we first ‘clicked’ a simple fluorescent moiety, Alexa Fluor 555-labeled DBCO (*i.e.*, DBCO-Alexa555), before testing the impact of adding more complex fluorescent signal generating probes (see further). Following the three labelling approaches (**Figure 1A-C**), the respective CRISPR RNP complexes were incubated with a 433-basepair (bp) dsDNA target, labeled with ATTO 488 (dsDNA-ATTO488) and loaded on a 1% agarose gel to perform a fEMSA (**Figure S2**). The dsDNA-ATTO488 target has a telomere-like repeat region (*i.e.*, TTAGGG repeats), to match the used telomere-specific crRNA (**Table S1**).

Fluorescence colocalization in the fEMSA experiments confirmed that chemical modification of the CRISPR-dCas9 complex can be carried out in any of the three stages of *in vitro* RNP assembly, without affecting its DNA binding potential (**Figure 2B**, lanes 3-5). Ternary complex formation between CRISPR RNP and dsDNA-ATTO488 resulted in a band that ran higher in the gel compared to free dsDNA-ATTO488 (**Figure 2B**, lane 1 ‘No CRISPR’), leading to colocalization of ATTO488 (cyan) and Alexa555 (magenta) fluorescence. This was comparable to the positive control used in this experiment, where a CRISPR-dCas9 complex having native, fluorescently labeled tracrRNA (*i.e.*, Alt-R tracrRNA-ATTO 550) was bound to the same dsDNA-ATTO488 target (**Figure 2B,** lane 2, ‘Alt-R CRISPR-ATTO550’). Furthermore, we assessed the specificity of SPAAC click-chemistry by using free DBCO-Alexa555 and commercially-available native Alt-R tracrRNA, which is devoid of 5’ azide modification (lane 6, ‘Alt-R CRISPR + DBCO-Alexa555’). This control demonstrated the ability of CRISPR to bind dsDNA-ATTO488, while the absence of azide-labeled tracrRNA hindered SPAAC click-chemistry and yielded unlabeled CRISPR RNP. The specificity of CRISPR-DNA ternary complex formation was shown using *GAL4* crRNA (lane 7, ‘Alt-R CRISPR*_GAL4_*-ATTO550**’**), which does not recognize dsDNA-ATTO488 target bearing a telomere-like repeat region. Finally, lanes 8 and 9 depict the free fluorescent gRNA (‘Only Alt-R gRNA-ATTO550**’**) and excess of unbound DBCO-Alexa555 (‘Only DBCO-Alexa555’). Collectively, these results show that (1) DBCO-fluorophore can be successfully ‘clicked’ (via SPAAC click-chemistry) to the three stages of RNP formation, as hypothesized earlier, without interfering with CRISPR complex formation, and (2) CRISPR binding functionality remains preserved as demonstrated by CRISPR-dsDNA complex formation.

### Fluorophore ‘clicked’ at three RNP formation stages preserves CRISPR capacity to bind genomic DNA *in situ*

After *in vitro* confirmation, we assessed the ability of the engineered CRISPR complexes to bind to genomic DNA in fixed U2OS cells. The fluorescent CRISPR-Alexa555 complexes were targeted towards human telomeres, given their widespread adoption as reference targets for developing CRISPR imaging strategies.

In line with the fEMSA observations, acquired images (**Figure 3A**) demonstrated telomere visualization in fixed cells by engineered CRISPR complexes for each of the three labelling approaches. The median fluorescent telomere foci count per nucleus of the imaged cells was 68.0, 67.0, and 68.0 for the ‘tracrRNA-click’, ‘gRNA-click’ and ‘RNP-click’, respectively (**Figure 3B and C**). The foci count distributions, statistically not different among the three tested conditions (Kruskal-Wallis test, p > 0.05), reflected the heterogeneity of the chromosomal states of U2OS cells. To confirm that the fluorescent foci are indeed telomere regions, CRISPR-Alexa555 (tracrRNA-click) fluorescence imaging was performed simultaneously with PNA-FISH. Concretely, while CRISPR complexes were targeted towards the CCCTAA telomere repeats, PNA-FISH probes were designed to hybridize to the complementary TTAGGG strand. As expected, telomere-specific signal generation was confirmed by colocalized CRISPR and PNA-FISH fluorescence (**Figure 3D**), while non-specific fluorescence cross-talk of Alexa555-labeled CRISPR in the FITC channel (*i.e.*, PNA-FISH readout channel) was ruled out (**Figure S3**).

**Figure 3.**
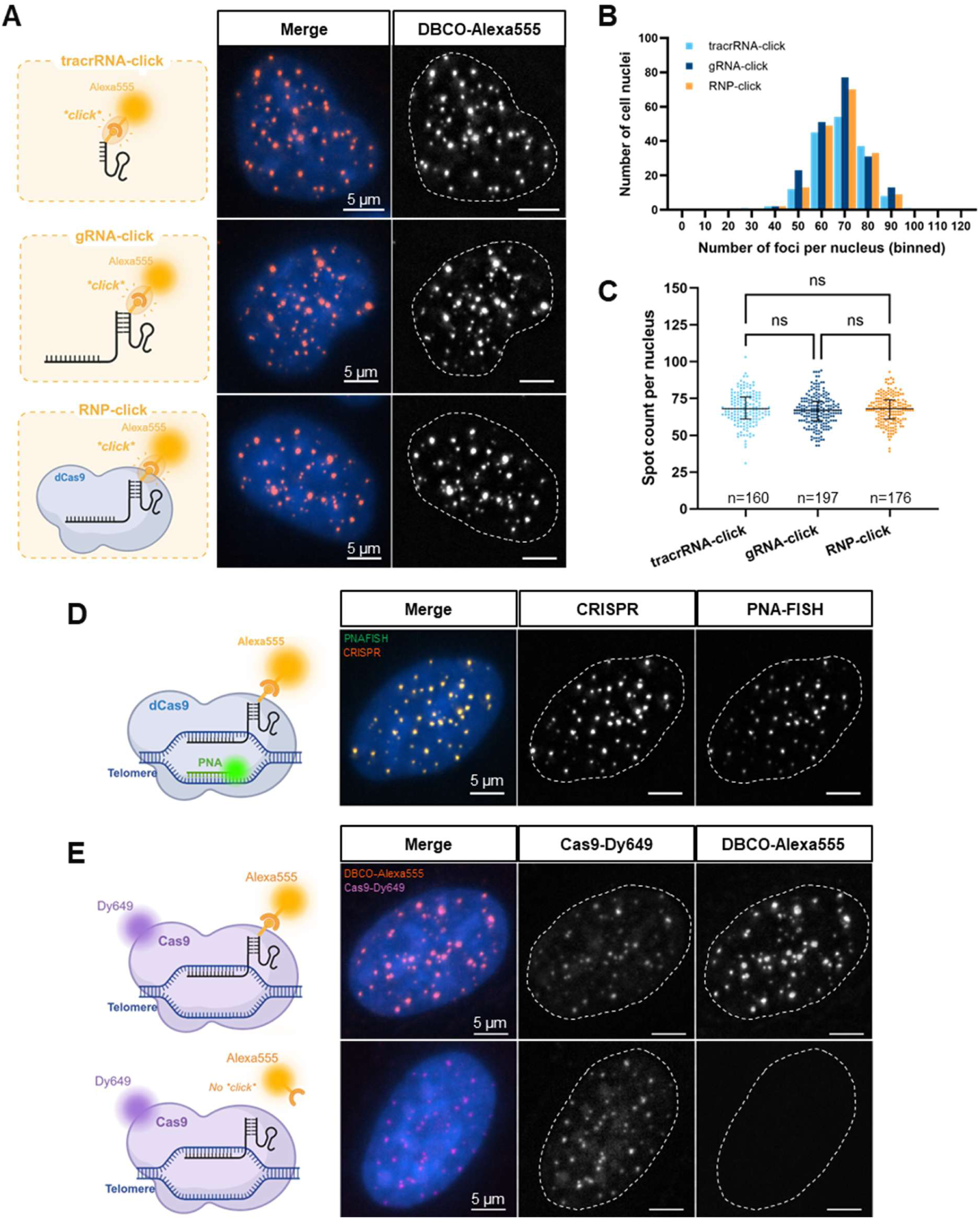
Fluorophore SPAAC click-chemistry preserves CRISPR capacity to bind telomere DNA in fixed U2OS cells regardless of the RNP formation step. **(A)** Telomere-targeting CRISPR complexes, modified by SPAAC click-chemistry with DBCO-Alexa555, according to the three engineering methodologies (Figure 1A-C), were delivered to fixed U2OS cells. For each of the three engineering methodologies (i.e., tracrRNA-click, gRNA-click, and RNP-click), the nucleus depicted is representative of the entire population of cells imaged in that condition. The ‘Merge’ panels depict a merged view of the nuclear DAPI staining (blue) and DBCO-Alexa555-labeled CRISPR complexes (orange). **(B)** Histogram shows the fluorescent Alexa555 foci counted per nucleus for each of the three engineering methodologies, while scatter plot **(C)** shows the distribution of fluorescent Alexa555 foci counted per nucleus for each engineering methodology, with median and interquartile range in black. Kruskal-Wallis test with Dunn’s multiple comparisons post-hoc test was performed: ns (p>0.05). For each condition, n represents the number of analyzed nuclei of three technical replicates. **(D)** Merge panel shows yellow fluorescence by colocalization of CRISPR (orange) and PNA-FISH (green) labelling. **(E)** Top and bottom panel show tracrRNA-click engineering with tracrRNA5’Azide and native Alt-R tracrRNA, respectively, with DBCO-Alexa555. Functional CRISPR complexes were formed with fluorescent Cas9-Dy649 for validating nuclear localization on telomere DNA.

Further, we also assessed the specificity of the SPAAC click-chemistry itself. Following approach **Figure 1A** (*i.e.,* the ‘tracrRNA-click’), DBCO-Alexa555 was incubated with either tracrRNA-5’azide or molecule not suitable for SPAAC click-chemistry (*i.e.,* native Alt-R tracrRNA lacking 5’ azide modification). Subsequently, a functional CRISPR RNP was formed with crRNA and fluorescent Cas9-SNAP fusion labeled with Dy649 (see Materials and Methods and **Figure S4**), the latter essential to locate all Cas9 binding. Of note, only in this particular experiment we used in-house produced fluorescent Cas9. By telomere imaging in fixed U2OS cells, we demonstrated that Cas9 colocalizes with Alexa555-labeled tracrRNA (**Figure 3E**). Conversely, the absence of fluorescence when 5’ azide modification was not included, confirmed that SPAAC click-chemistry was necessary for tracrRNA labelling and underlines the specificity of the reaction. Finally, as negative control, the CRISPR complexes were targeted towards the *GAL4* gene, which is absent in the human genome, and showed no significant binding to the fixed cells (**Figure S5**). Collectively, these results confirm the bio-orthogonal specificity of SPAAC click-chemistry and its capacity to robustly label CRISPR complexes for imaging applications.

### NA SPAAC click-chemistry preserves CRISPR capacity to bind genomic DNA

To enable broad applicability of the SPAAC click-chemistry, we next assessed whether it could allow the CRISPR complex to be labeled with other molecules, such as a DNA oligonucleotide. In contrast to a fluorophore, a DNA oligonucleotide can serve as a basis for advanced downstream signal generation strategies (*e.g.*, hybridization chain reaction, RCA). To this aim, a 27-nucleotide (nt) DNA oligonucleotide (with arbitrary sequence, **Table S1**), bearing DBCO molecule with a hydrophilic triethyleneglycol (TEG) linker (*i.e.*, DBCO-TEG, **Figure S6**) was utilized. Here, the TEG linker has a dual role: it increases solubility of the DBCO-DNA molecule while also providing more steric freedom for the DNA molecule to interact with the azide-modified tracrRNA in any of the three engineering methodologies (**Figure 1A-C**). Analogous to the fluorophore, the DBCO-labeled fluorescent oligonucleotide (*i.e.*, DBCO-TEG-DNA-ATTO550) was ‘clicked’ through SPAAC click-chemistry at the three CRISPR RNP formation stages. This comparison allowed us to investigate variations in SPAAC click-chemistry effectiveness caused by, for instance, non-specific interactions with the tracrRNA or gRNA itself, or steric hindrance of the ‘clicked’ DNA-ATTO550 interfering with gRNA-dCas9 recognition. *In vitro* fEMSA demonstrated successful modification of CRISPR complexes with DBCO-TEG-DNA-ATTO550 at all three stages of RNP formation, and their retained capacity to bind dsDNA-ATTO488 (**Figure 4,** lane 3, 4, and 5). Interestingly, CRISPR complexes formed after the ‘tracrRNA-click’ and ‘gRNA-click’ reactions showed decreased capacity to bind dsDNA compared to the ‘RNP’-click reaction, which suggests that such ‘click’ workflows interfere with CRISPR complex formation or dsDNA binding capacity. Lane 6 depicts the specificity of the SPAAC click-chemistry reaction in absence of an azide moiety, while lane 7 shows CRISPR binding specificity when *GAL4* crRNA was used. Finally, lanes 8 and 9 served to respectively reference the location of free gRNA and excess DBCO-TEG-DNA-ATTO550 in the gel.

**Figure 4.**
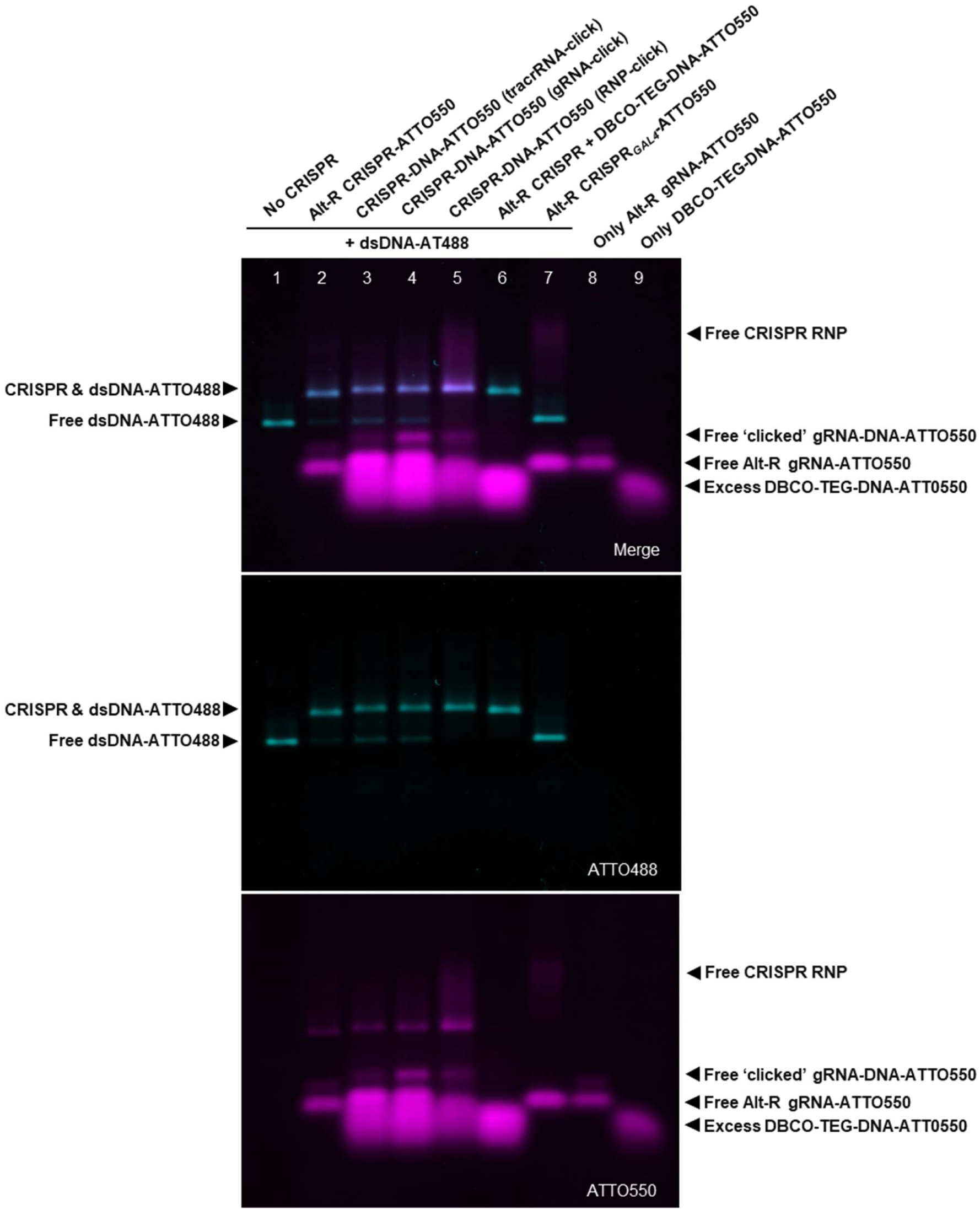
SPAAC click-chemistry enables CRISPR engineering with DNA at all three stages of RNP formation. fEMSA of DNA SPAAC-‘clicked’ CRISPR complexes and subsequent binding to dsDNA-ATTO488. The ‘Merge’ panel shows (colocalized) fluorescence by dsDNA-ATTO488 (cyan) and ATTO550 fluorescence through CRISPR-ATTO550 or DBCO-TEG-DNA-ATTO550 ‘clicked’ CRISPR complexes (magenta). Lane 1: reference dsDNA-ATTO488 in absence of CRISPR complex (‘No CRISPR’). Lane 2: a positive control showing native CRISPR complex, with fluorescently labeled Alt-R tracrRNA-ATTO550, binding the dsDNA-ATTO488 (‘Alt-R CRISPR-ATTO550’). Lanes 3-5: the SPAAC click reaction with DBCO-TEG-DNA-ATTO550 according to the three engineering methodologies as demonstrated in Figure 1 (‘CRISPR-DNA-ATTO550’). Lane 6: a native CRISPR complex with Alt-R tracrRNA, in absence of azide modification and in presence of DBCO-TEG-DNA-ATTO550, binding to dsDNA-ATTO488 (‘Alt-R CRISPR + DBCO-TEG-DNA-ATTO550’). Lane 7: GAL4-targeted CRISPR complex with Alt-R tracrRNA-ATTO550 that is unable to bind dsDNA-ATTO488 (‘Alt-R CRISPR_GAL4_-ATTO550’). Lanes 8-9: free fluorescent gRNA with Alt-R tracrRNA-ATTO550 (‘Only Alt-R gRNA-ATTO550’) and excess DBCO-TEG-DNA-ATTO550 (‘Only DBCO-TEG-DNA- ATTO550’), respectively.

Subsequently, the engineered CRISPR complexes were assessed for their ability to image telomeres in fixed U2OS cells. In addition, we investigated whether the fluorescent DNA oligonucleotide ‘clicked’ to the CRISPR complex would remain accessible in situ to enable more complex signal generation strategies (see also next section). Access to the fluorescent DNA was determined by hybridization of a complementary fluorescent probe. Therefore, we first delivered the DNA-ATTO550-modified CRISPR complexes to fixed U2OS cells (**Figure 5A, middle column**), followed by hybridization of a secondary ATTO 488-labeled DNA hybridization probe (DNAHyb-ATTO488) for fluorescence colocalization (**Figure 5A, right column**).

**Figure 5.**
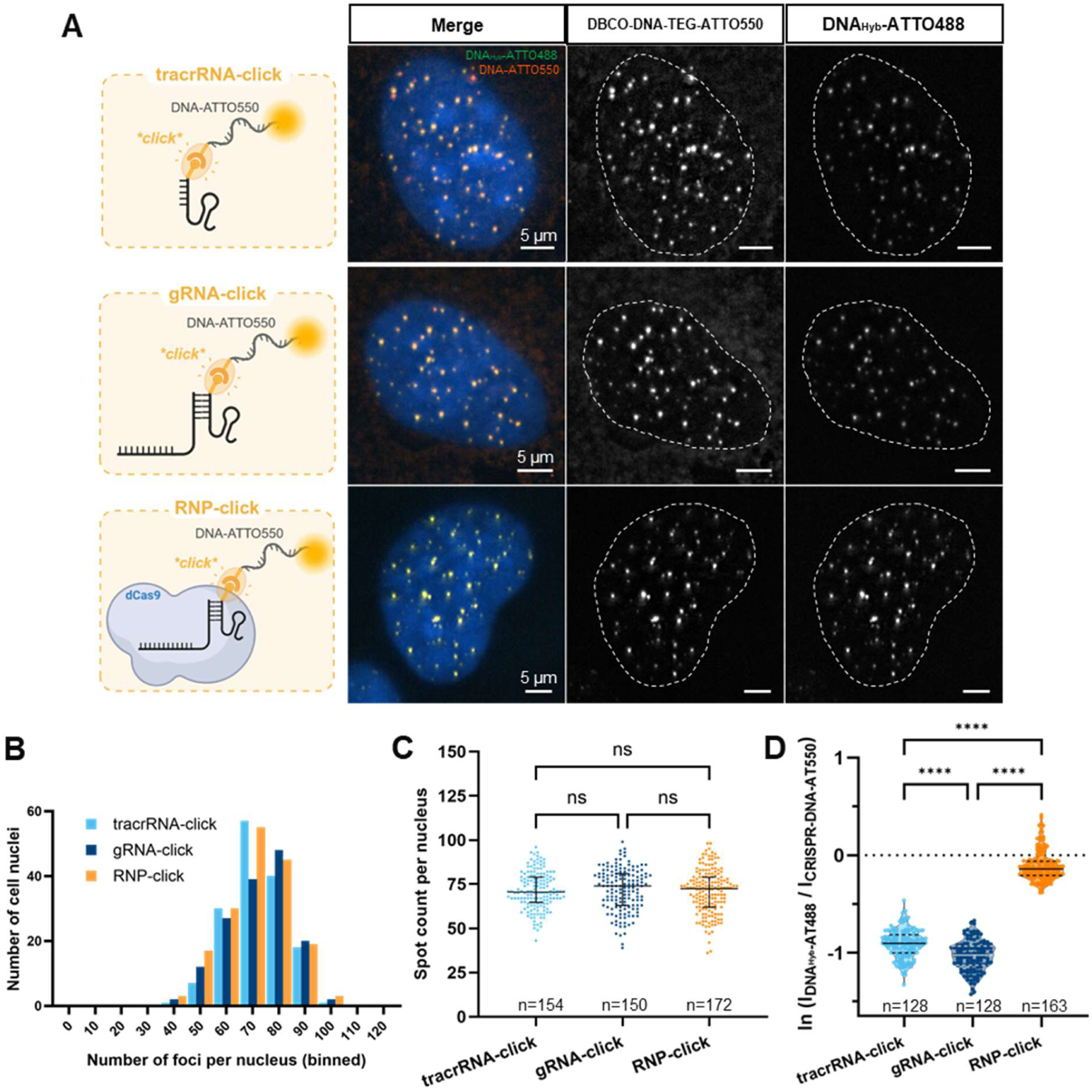
NA SPAAC click-chemistry preserves CRISPR capacity to bind genomic DNA and remains accessible for probe hybridization-based signal generation. **(A)** Telomere-targeting CRISPR complexes, modified by SPAAC click-chemistry with DBCO-TEG-DNA-ATTO550 according to the three engineering methodologies (Figure 1A-C), were delivered to fixed U2OS cells. For each of the engineering methodologies (i.e., tracrRNA-click, gRNA-click, RNP-click), the nucleus depicted is representative of the entire population of cells imaged in that condition. The ‘Merge’ panels depict a merged view of the nuclear DAPI stain (blue), DBCO-TEG-DNA-ATTO550-labeled CRISPR complexes (orange), and secondary ATTO488-labeled DNA hybridization probe (DNAHyb-ATTO488, green) colocalizing as yellow foci. **(B)** Histogram displays the fluorescent CRISPR-DNA-ATTO550 foci counted per nucleus for each of the three engineering methodologies, while scatter plot **(C)** shows the distribution of fluorescent CRISPR-DNA-ATTO550 foci counted per nucleus for each of the engineering methodologies with median and interquartile range in black. Kruskal-Wallis test with Dunn’s multiple comparisons post-hoc test was performed: ns (p>0.05), **** (p< 0.0001). n is the number of analyzed nuclei from three technical repetitions. **(D)** The hybridization efficiency ratio per nucleus was calculated by dividing the average DNAHyb-ATTO488 intensity (IDNAHyb-ATTO488) by the average CRISPR-DNA-ATTO550 intensity (ICRISPR-DNA-ATTO550) for each nucleus. This ratio was logarithmically transformed (natural log) and is represented in the violin plot (each datapoint represents one nucleus) with the median (full line) and interquartile range (dotted line) shown. Ordinary one-way ANOVA with Tukey’s multiple comparisons post-hoc test was performed: **** (p< 0.0001). n is the number of analyzed nuclei from three technical repetitions.

Quantification of the number of fluorescent ATTO550 telomere foci per nucleus revealed statistically similar median values (Kruskal-Wallis, p < 0.05) among the three SPAAC click-chemistry approaches: 70.5, 74.0, and 72.5, for ‘tracrRNA-click’, ‘gRNA-click’, and ‘RNP-click’, respectively (**Figure 5B and C**). As anticipated, the heterogeneity of the chromosomal states of the cells that drives the foci count distribution per condition remained coherent among the three approaches (demonstrated by absence of significant differences in **Figure 5C** after Kruskal-Wallis test, p > 0.05).

We further investigated the hybridization of DNA_Hyb_-ATTO488 to the DNA-ATTO550-modified CRISPR complexes for the three SPAAC click-chemistry approaches. Here, the average foci intensity was extracted per nucleus for both CRISPR-DNA-ATTO550 and DNA_Hyb_-ATTO488 fluorescence and used for calculating a hybridization efficiency ratio (I_DNAHyb-ATTO488_ / I_CRISPR-DNA-ATTO550_), which was subsequently natural log-transformed and plotted in **Figure 5D**. Interestingly, this ratio was significantly higher for the ‘RNP-click’ strategy (compared to the ‘tracrRNA-click’ and ‘gRNA-click’), which was driven by a relatively lower fluorescence intensity contribution of CRISPR-DNA-ATTO550, and a relatively higher DNA_Hyb_-ATTO488 fluorescence intensity contribution (**Figure S7**). Consistent with earlier observations on perturbed CRISPR RNP’s DNA binding capacity for ‘tracrRNA-click’ and ‘gRNA-click’ (Figure 4), we hypothesize that in the ‘RNP-click’ approach, the ‘clicked’ DNA-ATTO550 does not interfere with RNP formation (as RNP assembly precedes the SPAAC click-chemistry with DNA-ATTO550). As such, the ‘clicked’ oligonucleotide remains more accessible for hybridization, however, at the cost of lower SPAAC ‘click’ efficiency due to steric hindrance by the dCas9 protein. In contrast, for ‘tracrRNA-click’ and ‘gRNA-click’, the click-chemistry reaction of DNA-ATTO550 with the tracrRNA precedes crRNA and dCas9 interactions during RNP assembly. SPAAC click-chemistry therefore appears more efficient due to a higher degree of steric freedom. Yet, RNP formation interactions might interfere with the accessibility of the ‘clicked’ DNA-ATTO550 for later hybridization with DNA_Hyb_-ATTO488, while the excitation-emission of the ATTO 550 fluorophore on its own remains relatively unaffected. Such events would favor CRISPR-DNA-ATTO550 fluorescence intensity and relatively penalize hybridization-driven DNA_Hyb_-ATTO488 fluorescence intensity for ‘tracrRNA-click’ and ‘gRNA-click’ methodology, and hence lead to a lower hybridization efficiency ratio compared to the ‘RNP-click’ engineering strategy. Collectively, these results demonstrate that DNA ‘clicked’ to the CRISPR complex through SPAAC click-chemistry remains accessible for hybridization, while favoring ‘RNP-click’ strategy over the other two engineering approaches.

### NA SPAAC enables novel CRISPR-RCA approach

Having established robust SPAAC click-chemistry for modifying CRISPR complexes with DNA, we further explored this engineering approach and its unique advantages to accommodate more complex signal generation strategies. Interestingly, the modular nature of SPAAC allows covalent linking of different NAs regardless of their 5’-3’ orientation, even linking RNA and DNA, as explored in this work. We leveraged this unique capability for coupling an RCA reaction to the bound CRISPR complex, as a signal amplification strategy. Concretely, we labeled the CRISPR complex for the first time with a DNA primer that binds to a dumbbell-shaped RCA template (**Figure S8A** and explanation further), previously described as a strategy to improve RCA specificity and to physically compact the RCA product (RCP) (38). The dumbbell RCA protocol was established *in vitro* (**Figure S8B, C**) before being adapted for CRISPR-based imaging in fixed cells (**Figure 6A**). The previously described insights demonstrate that the ‘RNP-click’ engineering strategy preserves accessibility of the ‘clicked’ DNA better than the other two SPAAC click-chemistry approaches. Therefore, the 20-mer DNA oligonucleotide with free 3’ end, functioning as RCA primer (**Table S1**), was ‘clicked’ to the 5’ azide modified tracrRNA according to the ‘RNP-click’ methodology. Subsequently, the modified complexes were delivered to fixed U2OS cells to bind telomeres, followed by primer-template hybridization to directly initiate RCA by Phi29 polymerase in presence of dNTPs. Finally, readout of the RCP was carried out by hybridization of ATTO 550-labeled DNA probes DNA_RCA_-ATTO550 (**Table S1**).

**Figure 6.**
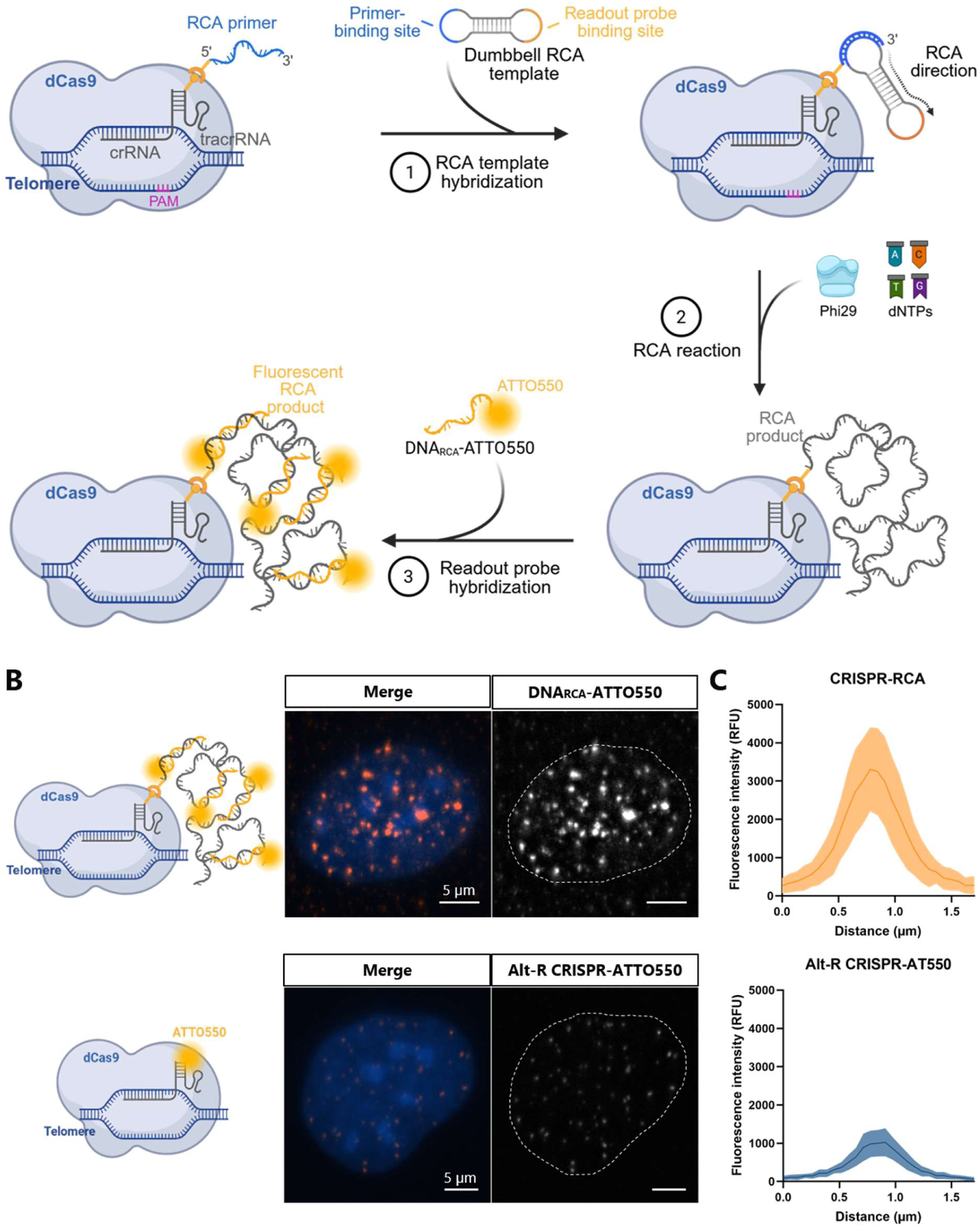
SPAAC engineering of CRISPR RNP allows RCA-based signal amplification. **(A)** First, the RCA primer was ‘clicked’ on the CRISPR complex by the ‘RNP-click’ engineering strategy, followed by delivering the complex to fixed U2OS cells. Subsequently, the RCA dumbbell template was delivered to the cells for hybridization to the CRISPR-RCA primer construct, followed by RCA amplification with Phi29 polymerase in presence of dNTPs. Ultimately, ATTO 550-labeled fluorescent readout probes (DNA_RCA_-ATTO550) were hybridized to the RCP for fluorescence readout. **(B)** Telomere imaging by the CRISPR-RCA approach (top) in comparison with the unamplified Alt-R CRISPR-ATTO550 complex (bottom). Image acquisition settings for this comparative experiment required lower exposure time such that RCA- amplified signal intensity could be captured within the dynamic range of the microscope, hence the observable difference in spot intensity and size for the reference Alt-R CRISPR-ATTO550 complex. **(C)** Plot displays the fluorescence intensity profile (average ± standard deviation) of telomere spots imaged through CRISPR-RCA-enabled signal amplification, compared to Alt-R CRISPR-ATTO550 imaging.

Telomere imaging was used to estimate the fluorescent signal gain resulting from RCA compared to native Alt-R CRISPR-ATTO550 (**Figure 6B**). Fluorescence intensity comparison underlined the level of amplification reached through the CRISPR-RCA methodology (**Figure 6C and D**): RCA increased signal strength by 400%, without significant spatial distortion. Simultaneously, absence of off-target amplification signal for negative control GAL4-targeted CRISPR-RCA and Alt-R CRISPR-ATTO550 based readout demonstrated the specificity of the reaction (**Figure S9**). Collectively, these results demonstrate how this new engineering methodology allows straightforward modification of the CRISPR complex for establishing new signal generations strategies for CRISPR-based genome imaging.

## DISCUSSION

CRISPR-based technologies have been increasingly implemented for genome imaging in fixed and live cells over the past years. That trend coincided with the development of various strategies for signal generation, and modalities such as signal amplification and multi-color readout. This typically required altering the CRISPR complex either by (d)Cas9 engineering or by implementing gRNA modifications. The latter has seen widespread adoption owing to higher flexibility of NA engineering compared to Cas proteins. However, gRNA engineering for including new functionalities in the CRISPR complex remains non-trivial. Previous studies showed that modifications to the gRNA have been implicated in reducing CRISPR performance *in vitro* and in imaging applications (20, 23, 42). For instance, modifications to the 3’-end of single-guide RNA ((s)gRNA) yielded poor imaging performance compared to internally-modified sgRNA, which was ascribed to interference with internal structures crucial for gRNA-(d)Cas9 recognition and complex formation (23). Mitigation strategies, such as the inclusion of linkers to spatially separate aptamers from crucial gRNA-Cas structures still showed gene target-dependent performance (20). These observations suggest that gRNA engineering success depends on the location and RNA-structural context of the implemented modification. Furthermore, while these dynamics remain poorly understood, there are no approaches that possess the flexibility to troubleshoot and circumvent these design challenges.

In this context, we utilized the versatility of SPAAC click-chemistry to enable highly controllable modification of the CRISPR RNP with the flexibility of decoupling it from native RNP assembly. Specifically, 5’ azide-modified tracrRNA was used as a nucleation site for SPAAC reactions to covalently link fluorophores or different DNA molecules, thereby demonstrating that these reactions can take place throughout the stages of functional CRISPR RNP formation (*i.e.*, either before or after gRNA annealing, or especially after (d)Cas9 RNP formation). This implies that with increasing steric strain and changing electrostatic interactions, going from isolated tracrRNA ultimately to RNP formation, the azide-modified tracrRNA remains available for conjugating DBCO-modified molecules. Further, our findings show that when SPAAC click-chemistry with DNA precedes dCas9-gRNA assembly, CRISPR-target recognition and the accessibility of the ‘clicked’ DNA are affected. Conversely, performing DNA SPAAC ‘click’ only after dCas9-gRNA assembly benefits CRISPR-target recognition and accessibility of the ‘clicked’ DNA. These insights are in line with the previously-described CRISPR engineering and functionality challenges in the genome imaging field, while also providing a framework to resolve them. Ultimately, the versatility of the engineering strategy was consolidated in a novel CRISPR-RCA approach for amplified signal generation. While other applications required internal gRNA modification and a layer of secondary and tertiary bridging probes to achieve the necessary scaffold for RCA (43, 44), our SPAAC-based approach circumvents such complex constructs, while attaching the RCP covalently to CRISPR. As such, we demonstrate how RNA-DNA hybrid NAs can be elegantly engineered with high control and possibility to overcome 5’-3’ directionality constraints, which both can facilitate new strategies for CRISPR applications.

Here we present a new framework to rationally engineer CRISPR complexes, while recognizing that there is still much to be gained from deeper mechanistic understanding. Our work has focused initially on fixed cells, which is relevant for integrating CRISPR-based genome imaging with spatial multi-omic imaging protocols applied in fixed cells and tissues (45). Demonstrating this framework in live-cell imaging context could potentially further prove its versatility. Concretely, the vast majority of CRISPR-based live-cell imaging technologies relies on plasmid-based or viral delivery for stable expression of engineered CRISPR components, such as aptamer-engineered (s)gRNAs, into living cells. Therefore, the design of these engineered sgRNA structures is confined by plasmid design constraints that do not permit the expression of non-native hybrid DNA-RNA NAs, while the natural expression processes of a living cell limit timely control over the engineering step. Furthermore, the continuous expression of CRISPR components in live cells is known to cause chromatin perturbations, and limits its use to immortalized cell lines (16, 46). In contrast, our framework goes beyond traditional in vitro NA synthesis or expression-based live-cell delivery by dissociating native in vitro RNP assembly from engineering step, minimizing structural perturbation of CRISPR and preserving downstream imaging functionality. As such, this engineering framework combined with a CRISPR RNP-based delivery approach opens up new possibilities for live-cell genome imaging studies either by resolving the engineering challenges of existing signal generation approaches or by developing entirely new ways to do so. If CRISPR-based imaging, similar to gene editing, is to become a gold standard fixed- and live-cell genome imaging technique for fundamental research and especially clinical diagnostics, such RNP-based engineering and delivery approaches will be crucial (47, 48). Ultimately, the proposed framework has applications beyond genome imaging. For example, the groundbreaking CRISPR-based prime editing technology relies on engineering the sgRNA extensively, for which it is known that self-hybridization, and steric and torsional hindrances interfere with Cas9-gRNA interaction and the gene editing performance thereof (3, 49). With the utilization of our engineering framework, gene editing efficiency gains are achievable.

## ACKNOWLEDGEMENTS

We would like to thank Prof. Dirk Daelemans for kindly providing the pLentiCRISPR-Cas9 construct.

## AUTHOR CONTRIBUTIONS

Charlotte Van Tricht: Conceptualization, Data curation, Formal analysis, Funding acquisition, Investigation and Methodology, Visualisation, Writing: original draft, Writing: review & editings. Saphyra Thonon: Formal analysis, Investigation and Methodology, Visualization, Writing: review & editing. David Wouters: Formal analysis, Investigation and Methodology, Software, Writing: review & editing. Alejandro Sifrim: Conceptualization, Funding acquisition, Writing: review & editing. Katy Vandereyken: Conceptualization, Funding acquisition. Vitor B. Pinheiro: Conceptualization, Funding acquisition, Writing: review & editing. Thierry Voet: Conceptualization, Funding acquisition, Writing: review & editing. Jeroen Lammertyn: Conceptualization, Funding acquisition, Writing: review & editing. Dragana Spasic: Conceptualization, Funding acquisition, Visualization, Writing: original draft, Writing: review & editing.

## SUPPLEMENTARY DATA

Supplementary Data are available at NAR online.

## DATA AVAILABILITY

The data underlying this article will be shared on reasonable request to the corresponding author.

## REFERENCES

1. Ran, F.A., Hsu, P.D., Wright, J., Agarwala, V., Scott, D.A. and Zhang, F. (2013) Genome engineering using the CRISPR-Cas9 system. Nat. Protoc., 8, 2281–2308.

2. Komor, A.C., Kim, Y.B., Packer, M.S., Zuris, J.A. and Liu, D.R. (2016) Programmable editing of a target base in genomic DNA without double-stranded DNA cleavage. Nature, 533, 420–424.

3. Anzalone, A. V., Randolph, P.B., Davis, J.R., Sousa, A.A., Koblan, L.W., Levy, J.M., Chen, P.J., Wilson, C., Newby, G.A., Raguram, A., et al. (2019) Search-and-replace genome editing without double-strand breaks or donor DNA. Nature, 576, 149–157.

4. Gilbert, L.A., Larson, M.H., Morsut, L., Liu, Z., Brar, G.A., Torres, S.E., Stern-Ginossar, N., Brandman, O., Whitehead, E.H., Doudna, J.A., et al. (2013) CRISPR-mediated modular RNA-guided regulation of transcription in eukaryotes. Cell, 154, 442.

5. Sander, J.D. and Joung, J.K. (2014) CRISPR-Cas systems for editing, regulating and targeting genomes. Nat. Biotechnol., 32.

6. Hajian, R., Balderston, S., Tran, T., deBoer, T., Etienne, J., Sandhu, M., Wauford, N.A., Chung, J.Y., Nokes, J., Athaiya, M., et al. (2019) Detection of unamplified target genes via CRISPR–Cas9 immobilized on a graphene field-effect transistor. *Nat*. Biomed. Eng., 3, 427–437.

7. Li, Y., Zhao, W., Wu, Y., Li, R., Zhang, J., Xie, H., Zhang, K. and Li, J. (2025) In vivo CRISPR biosensing. Chem. Soc. Rev., 10.1039/d5cs00921a.

8. Henssen, A.G. (2022) Shedding Light on ecDNA Behavior Using CRISPR-Based Live-Cell Imaging. Cancer Discov., 12, 293–295.

9. Van Tricht, C., Voet, T., Lammertyn, J. and Spasic, D. (2022) Imaging the unimaginable: leveraging signal generation of CRISPR-Cas for sensitive genome imaging. Trends Biotechnol., 41, 769–784.

10. Chen, B., Gilbert, L.A., Cimini, B.A., Schnitzbauer, J., Zhang, W., Li, G.W., Park, J., Blackburn, E.H., Weissman, J.S., Qi, L.S., et al. (2013) Dynamic imaging of genomic loci in living human cells by an optimized CRISPR/Cas system. Cell, 155, 1479–1491.

11. Ma, H., Naseri, A., Reyes-Gutierrez, P., Wolfe, S.A., Zhang, S. and Pederson, T. (2015) Multicolor CRISPR labeling of chromosomal loci in human cells. Proc. Natl. Acad. Sci. U. S. A., 112, 3002–3007.

12. Chen, B., Hu, J., Almeida, R., Liu, H., Balakrishnan, S., Covill-Cooke, C., Lim, W.A. and Huang, B. (2016) Expanding the CRISPR imaging toolset with Staphylococcus aureus Cas9 for simultaneous imaging of multiple genomic loci. Nucleic Acids Res., 44, e75.

13. Xu, H., Wang, J., Liang, Y., Fu, Y., Li, S., Huang, J., Xu, H., Zou, W. and Chen, B. (2020) TriTag: an integrative tool to correlate chromatin dynamics and gene expression in living cells. Nucleic Acids Res., 48, e127.

14. Tanenbaum, M.E., Gilbert, L.A., Qi, L.S., Weissman, J.S. and Vale, R.D. (2014) A protein-tagging system for signal amplification in gene expression and fluorescence imaging. Cell, 159, 635–646.

15. Ye, H., Rong, Z. and Lin, Y. (2017) Live cell imaging of genomic loci using dCas9-SunTag system and a bright fluorescent protein. Protein Cell, 8, 853–855.

16. Hou, M., Li, Y., Wu,· Xiaobo, Long,· Die, Di Sun, Chen, Peizhen and Huang, H. (2026) Recent Advances in the Development of CRISPR-Based Live-Cell Molecular Imaging and Sensing. Mol. Imaging Biol., 10.1007/S11307-026-02110-Y.

17. Qin, P., Parlak, M., Kuscu, C., Bandaria, J., Mir, M., Szlachta, K., Singh, R., Darzacq, X., Yildiz, A. and Adli, M. (2017) Live cell imaging of low- and non-repetitive chromosome loci using CRISPR-Cas9. Nat. Commun., 8, 1–10.

18. Lyu, X.Y., Deng, Y., Huang, X.Y., Li, Z.Z., Fang, G.Q., Yang, D., Wang, F.L., Kang, W., Shen, E.Z. and Song, C.Q. (2022) CRISPR FISHer enables high-sensitivity imaging of nonrepetitive DNA in living cells through phase separation-mediated signal amplification. Cell Res., 32, 969–981.

19. Clow, P.A., Du, M., Jillette, N., Taghbalout, A., Zhu, J.J. and Cheng, A.W. (2022) CRISPR-mediated multiplexed live cell imaging of nonrepetitive genomic loci with one guide RNA per locus. Nat. Commun., 13, 1871.

20. Yang, L.Z., Min, Y.H., Liu, Y.X., Gao, B.Q., Liu, X.Q., Huang, Y., Wang, H., Yang, L., Liu, Z.J. and Chen, L.L. (2024) CRISPR-array-mediated imaging of non-repetitive and multiplex genomic loci in living cells. Nature Methods 2024 21:9, 21, 1646–1657.

21. Cheng, A.W., Jillette, N., Lee, P., Plaskon, D., Fujiwara, Y., Wang, W., Taghbalout, A. and Wang, H. (2016) Casilio: A versatile CRISPR-Cas9-Pumilio hybrid for gene regulation and genomic labeling. Cell Res., 26, 254–257.

22. Ma, H., Tu, L.C., Naseri, A., Huisman, M., Zhang, S., Grunwald, D. and Pederson, T. (2016) Multiplexed labeling of genomic loci with dCas9 and engineered sgRNAs using CRISPRainbow. Nat. Biotechnol., 34, 528–530.

23. Ma, H., Tu, L.C., Naseri, A., Chung, Y.C., Grunwald, D., Zhang, S. and Pederson, T. (2018) CRISPR-Sirius: RNA scaffolds for signal amplification in genome imaging. Nat. Methods, 15, 928–931.

24. Hong, Y., Lu, G., Duan, J., Liu, W. and Zhang, Y. (2018) Comparison and optimization of CRISPR/dCas9/gRNA genome-labeling systems for live cell imaging. Genome Biol., 19, 39.

25. Deng, W., Shi, X., Tjian, R., Lionnet, T. and Singer, R.H. (2015) CASFISH: CRISPR/Cas9-mediated in situ labeling of genomic loci in fixed cells. Proc. Natl. Acad. Sci. U. S. A., 112, 11870–11875.

26. Ma, Y., Wang, M., Li, W., Zhang, Z., Zhang, X., Wu, G., Tan, T., Cui, Z. and Zhang, X.E. (2017) Live Visualization of HIV-1 Proviral DNA Using a Dual-Color-Labeled CRISPR System. Anal. Chem., 89, 12896–12901.

27. Geng, Y. and Pertsinidis, A. (2021) Simple and versatile imaging of genomic loci in live mammalian cells and early pre-implantation embryos using CAS-LiveFISH. Sci. Rep., 11, 12220.

28. Zhang, S. and Song, Z. (2017) Aio-Casilio: a robust CRISPR–Cas9–Pumilio system for chromosome labeling. J. Mol. Histol., 48, 293–299.

29. Neguembor, M. V., Sebastian-Perez, R., Aulicino, F., Gomez-Garcia, P.A., Cosma, M.P. and Lakadamyali, M. (2018) (Po)STAC (Polycistronic SunTAg modified CRISPR) enables live-cell and fixed-cell super-resolution imaging of multiple genes. Nucleic Acids Res., 46, e30.

30. Mao, S., Wu, R., Luo, W., Qin, J. and Chen, A.K. (2025) Spuriously transcribed RNAs from CRISPR-sgRNA expression plasmids scaffold biomolecular condensate formation and hamper accurate genomic imaging. Nucleic Acids Res., 53, 192.

31. Agard, N.J., Prescher, J.A. and Bertozzi, C.R. (2004) A strain-promoted [3 + 2] azide-alkyne cycloaddition for covalent modification of biomolecules in living systems. J. Am. Chem. Soc., 126, 15046–15047.

32. Zou, R.S., Liu, Y. and Ha, T. (2021) In vitro Cleavage and Electrophoretic Mobility Shift Assays for Very Fast CRISPR. Bio. Protoc., 11, e4138.

33. Liu, Y., Zou, R.S., He, S., Nihongaki, Y., Li, X., Razavi, S., Wu, B. and Ha, T. (2020) Very fast CRISPR on demand. Science (1979)., 368, 1265–1269.

34. Taemaitree, L., Shivalingam, A., El-Sagheer, A.H. and Brown, T. (2019) An artificial triazole backbone linkage provides a split-and-click strategy to bioactive chemically modified CRISPR sgRNA. Nature Communications 2019 10:1, 10, 1–8.

35. Neggers, J.E., Vercruysse, T., Jacquemyn, M., Vanstreels, E., Baloglu, E., Shacham, S., Crochiere, M., Landesman, Y. and Daelemans, D. (2015) Identifying Drug-Target Selectivity of Small-Molecule CRM1/XPO1 Inhibitors by CRISPR/Cas9 Genome Editing. Chem. Biol., 22, 107–116.

36. Csibra, E., Renders, M. and Pinheiro, V.B. (2020) Bacterial Cell Display as a Robust and Versatile Platform for Engineering Low-Affinity Ligands and Enzymes. ChemBioChem, 21, 2844–2853.

37. Binder, U., Matschiner, G., Theobald, I. and Skerra, A. (2010) High-throughput Sorting of an Anticalin Library via EspP-mediated Functional Display on the Escherichia coli Cell Surface. J. Mol. Biol., 400, 783–802.

38. Wochner, A., Attwater, J., Coulson, A. and Holliger, P. (2011) Ribozyme-catalyzed transcription of an active ribozyme. Science (1979)., 332, 209–212.

39. Stringer, C., Wang, T., Michaelos, M. and Pachitariu, M. (2020) Cellpose: a generalist algorithm for cellular segmentation. Nature Methods 2020 18:1, 18, 100–106.

40. Van Der Walt, S., Schönberger, J.L., Nunez-Iglesias, J., Boulogne, F., Warner, J.D., Yager, N., Gouillart, E. and Yu, T. (2014) Scikit-image: Image processing in python. PeerJ, 2014, e453.

41. Nishimasu, H., Ran, F.A., Hsu, P.D., Konermann, S., Shehata, S.I., Dohmae, N., Ishitani, R., Zhang, F. and Nureki, O. (2014) Crystal structure of Cas9 in complex with guide RNA and target DNA. Cell, 156, 935–949.

42. George, J.T., Azhar, M., Aich, M., Sinha, D., Ambi, U.B., Maiti, S., Chakraborty, D. and Srivatsan, S.G. (2020) Terminal Uridylyl Transferase Mediated Site-Directed Access to Clickable Chromatin Employing CRISPR-dCas9. J. Am. Chem. Soc., 142, 13954–13965.

43. Zhang, K., Deng, R., Teng, X., Li, Y., Sun, Y., Ren, X. and Li, J. (2018) Direct Visualization of Single-Nucleotide Variation in mtDNA Using a CRISPR/Cas9-Mediated Proximity Ligation Assay. J. Am. Chem. Soc., 140, 11293–11301.

44. Liang, Y., Wu, S., Han, W., Wang, J., Xu, C., Shi, J., Zhang, Z., Gao, H., Zhang, K. and Li, J. (2022) Visualizing Single-Nucleotide Variations in a Nuclear Genome Using Colocalization of Dual-Engineered CRISPR Probes. Anal. Chem., 10.1021/acs.analchem.2c01208.

45. Rybczynski, A.T., Cottle, W.T., Chen, P.T., Kwon, J., Shang, T., Wang, Y., Meneses, P., Pangeni, S., Park, Y., Gavrilov, M., et al. (2025) Imaging the time course of DNA damage response at a nonrepetitive endogenous locus. Cell Rep. Methods, 5, 101219.

46. Xiao, Z. and Sun, Y. (2026) Illuminating the genome: emerging approaches in CRISPR-Cas live-cell imaging. Nucleic Acids Res., 54, 1540.

47. Karp, H., Zoltek, M., Wasko, K., Vazquez, A.L., Brim, J., Ngo, W., Schepartz, A. and Doudna, J.A. (2025) Packaged delivery of CRISPR–Cas9 ribonucleoproteins accelerates genome editing. Nucleic Acids Res., 53, 105.

48. Zhu, Y., Balaji, A., Han, M., Andronov, L., Roy, A.R., Wei, Z., Chen, C., Miles, L., Cai, S., Gu, Z., et al. (2025) High-resolution dynamic imaging of chromatin DNA communication using Oligo-LiveFISH. Cell, 188, 3310–3328.e27.

49. Chen, P.J. and Liu, D.R. (2022) Prime editing for precise and highly versatile genome manipulation. Nat. Rev. Genet., 24, 161–177.

