## Supplementary data for "MODULAR CLICK-CHEMISTRY-BASED ENGINEERING MINIMALLY PERTURBS CRISPR RIBONUCLEOPROTEIN FORMATION FOR GENOME IMAGING APPLICATIONS"

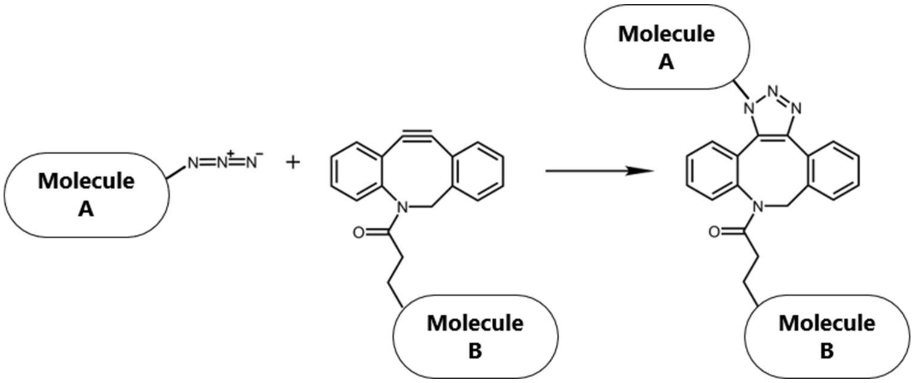


**Figure S1. Schematic of the bio-orthogonal SPAAC click-chemistry reaction.** Molecule A is modified with an azide functional group (*i.e.*, in this work tracrRNA$^{5'azide}$), while molecule B is functionalized with a DBCO functional group (*i.e.*, in this work DBCO-Alexa555, DBCO-TEG-DNA-ATTO550, DBCO-TEG-RCA primer). Upon reaction of these two functional groups, a triazole linkage is formed between molecules A and B through [3+2] cycloaddition. As such, two biomolecules can be covalently linked to each other.

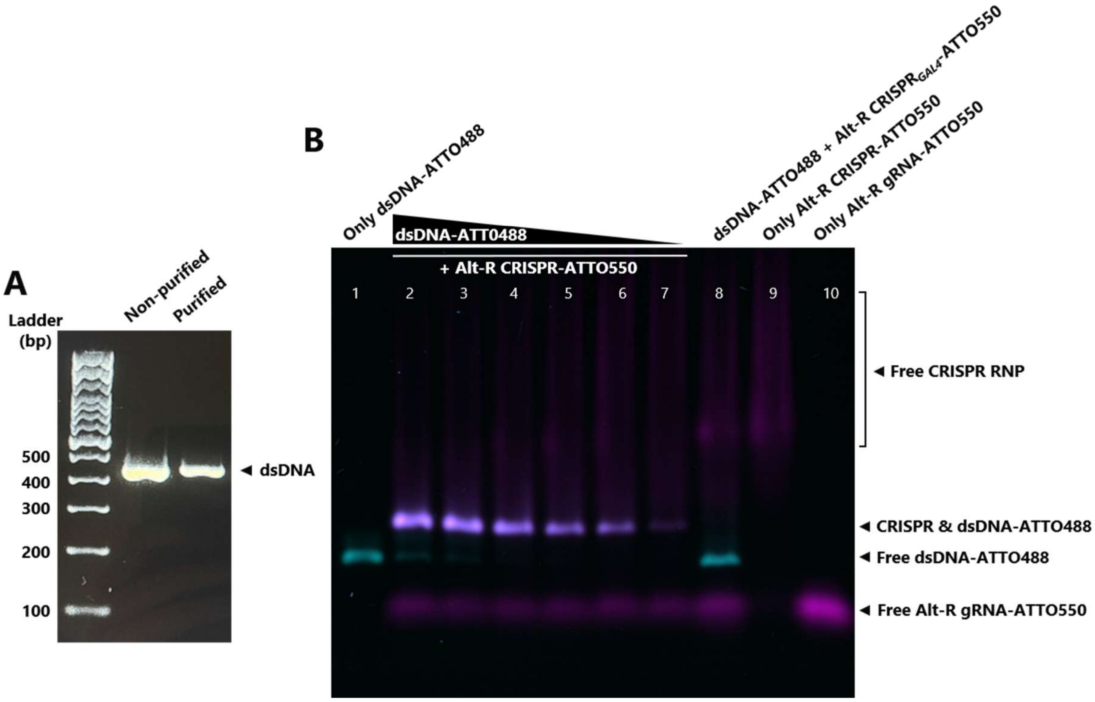


**Figure S2. PCR amplification and purification of dsDNA(-ATTO488), and its subsequent titration for fluorescent CRISPR-dCas9 fEMSA. (A)** After PCR amplification of the unlabeled dsDNA (433 bp), unpurified and purified samples were loaded on a 1% agarose gel and successful PCR amplification was confirmed against the ladder. **(B)** After performing

PCR with ATTO488-labeled forward primer, the resulting purified dsDNA-ATTO488 was titrated with a constant amount of native Alt-R CRISPR-ATTO550 complex, in order to determine the optimal amount of dsDNA-ATTO488 required for the fEMSA. For lanes 2-7, purified dsDNA-ATTO488 was added in the following concentrations: 5, 4, 3, 2, 1 and 0.5 µL, respectively.

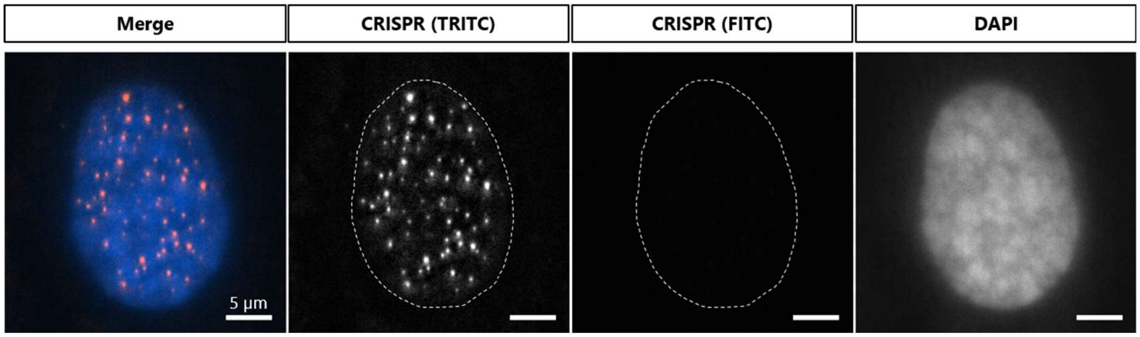


**Figure S3. Fluorescent signal generation by CRISPR-AF555 (tracrRNA-click) is specific, without fluorescent cross-talk in the FITC channel.** Telomere imaging with fluorescent CRISPR-Alexa555 complexes, engineered through the tracrRNA-click approach. No Alexa555 fluorescent cross-talk is captured in the FITC channel. Therefore, fluorescence in the FITC channel in Figure 3D is a result of ATTO488 labeled PNA probes only.

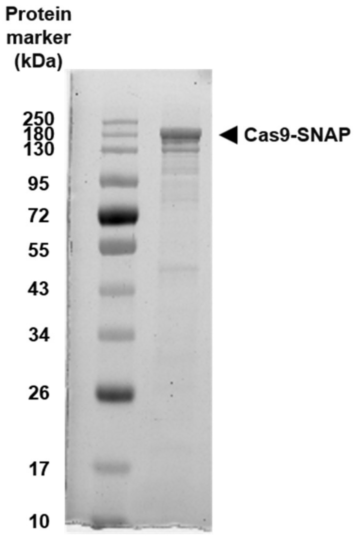


**Figure S4: SDS-PAGE analysis of purified Cas9-SNAP protein.** Loading 10 µL of a 1:10 dilution of the double purified Cas9-SNAP onto a 12% SDS-PAGE showed highly concentrated and purified Cas9-SNAP protein.

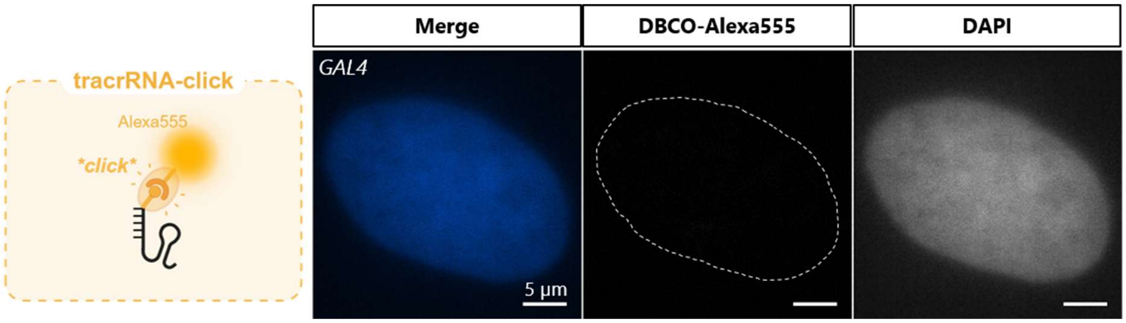


**Figure S5. Fluorescent signal generation by SPAAC click-chemistry-modified CRISPR$_{GAL4}$-Alexa555 complexes in fixed U2OS cells.** CRISPR complexes were modified with DBCO-Alexa555 by the 'tracrRNA-click' methodology, and targeted towards the GAL4 gene by an appropriate crRNA design. Given that the GAL4 gene is absent in the human genome, subsequent addition of these CRISPR$_{GAL4}$-Alexa555 complexes to fixed U2OS cells served as negative control and confirmed the specificity of CRISPR-based signal generation. The 'Merge' panel depicts a merged view of the nuclear DAPI staining (blue) and CRISPR$_{GAL4}$-Alexa555 complexes (orange). Schematics created with BioRender.com.

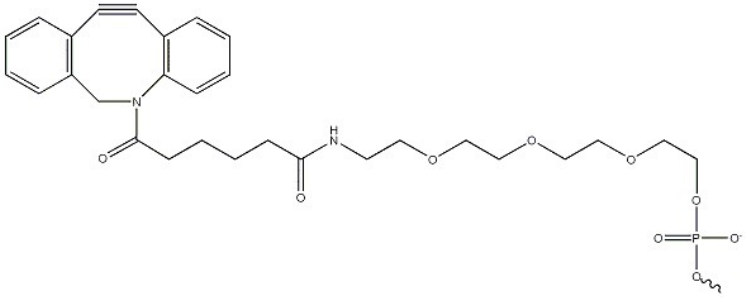

**Figure S6. DBCO with a TEG linker.** The TEG linker improves solubility of the entire molecule by its hydrophilic nature and allows steric freedom for the DNA oligonucleotide attached to it.

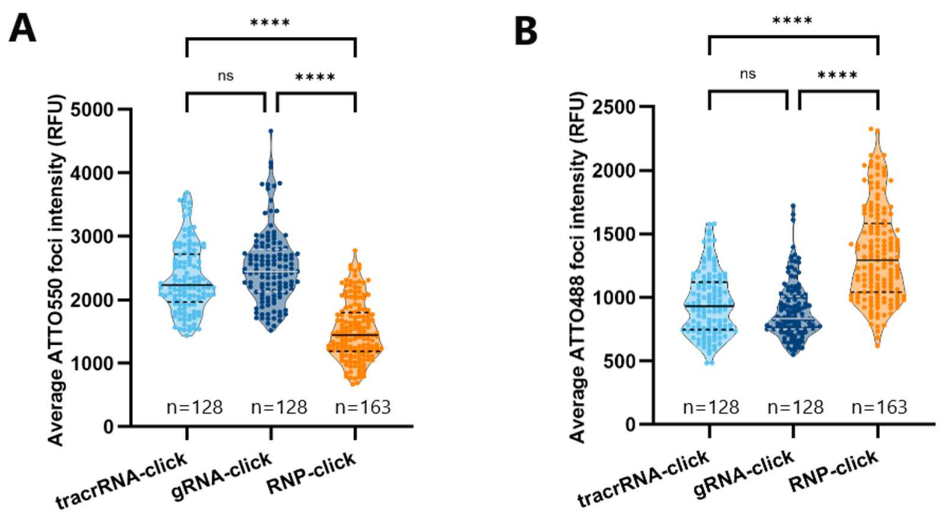


**Figure S7. Average intensity of CRISPR-DNA-ATTO550 and $DNA_{Hyb}$-ATTO488 foci for the 'tracrRNA-click', 'gRNA-click' and 'RNP-click' engineering strategies.** For each of the three engineering strategies, the average fluorescent foci intensity per nucleus for CRISPR-DNA-ATTO550 **(A)** and $DNA_{Hyb}$-ATTO488 **(B)** is shown in a violin plot with median (full line) and interquartile range (dotted line). Kruskal Wallis test with Dunn's multiple comparisons post-hoc test was performed: ns ($p>0.05$), **** ($p< 0.0001$). n is the number of nuclei from three technical repetitions.

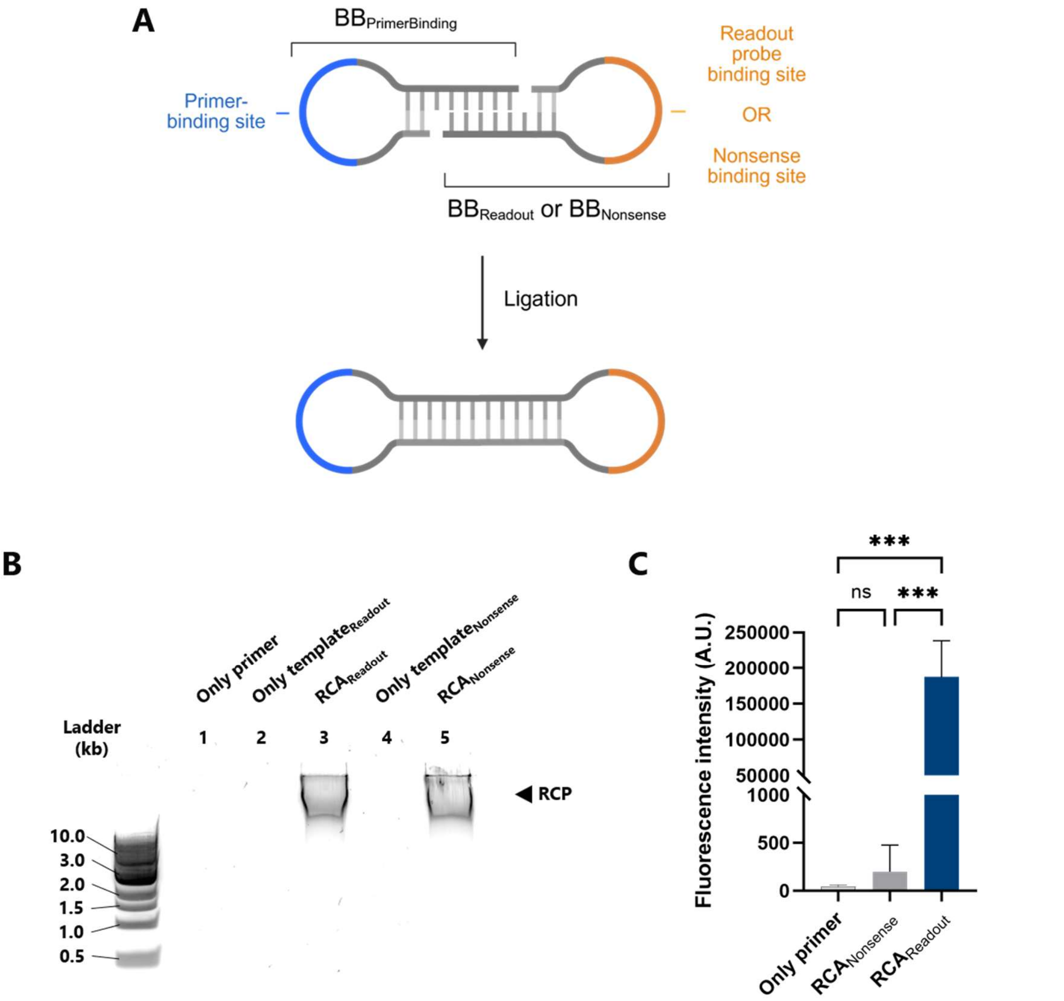


**Figure S8. In vitro development of dumbbell-templated RCA. (A)** Schematic representation of dumbbell RCA template construct. In order to generate the dumbbell template, two stem-loop building blocks $BB_{PrimerBinding}$ and $BB_{Readout}$ or $BB_{Nonsense}$ are annealed and ligated. Once ligated, the dumbbell RCA template can be used for RCA reactions in vitro or in situ. **(B)** Prior to adaption for CRISPR-based dumbbell-templated RCA, the standalone version of RCA (i.e., without SPAAC click-chemistry or CRISPR) was established in vitro. The in vitro RCP was loaded onto a 1% agarose gel after purification. Results show that RCA reactions were efficient for both the RCA template with readout probe binding site ('$RCA_{Readout}$', lane 3) and with nonsense binding site ('$RCA_{Nonsense}$', lane 5) and that reactions were template-dependent ('Only primer', lane 1) and primer-dependent ('Only $template_{Readout}$', lane 2  and 'Only $template_{Nonsense}$', lane 4). **(C)** End-point fluorescence intensity measurements demonstrated the fluorescent signal generation by hybridization of fluorescein-labeled readout probes to the in vitro RCP ('$RCA_{Readout}$'). The specificity of the assay was assessed by two negative control conditions ('Only primer', and '$RCA_{Nonsense}$'), where $RCA_{Nonsense}$ represents fluorescent signal generation, or the absence thereof, when RCA occurs with the $template_{Nonsense}$. Ordinary one-way ANOVA with Tukey's multiple comparisons was performed, with three technical repetitions per condition: ns ($p>0.05$), *** ($p< 0.0005$).

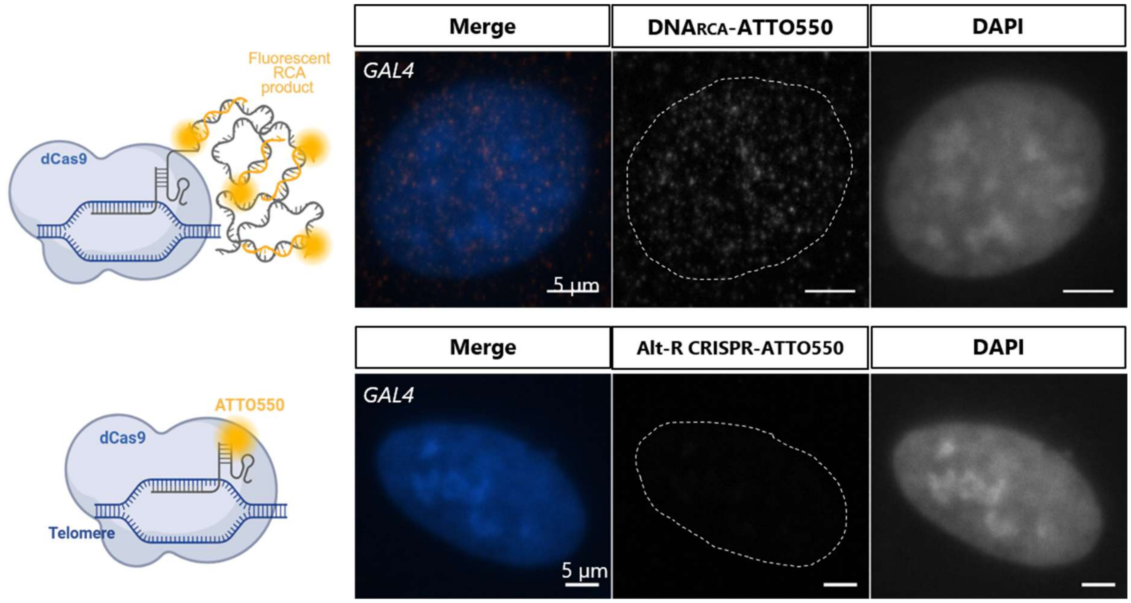


**Figure S9. GAL4 imaging by CRISPR-RCA and native Alt-R CRISPR-ATTO550 to assess fluorescent signal generation specificity.** GAL4, a yeast gene absent in the human genome, was imaged as negative control using identical image acquisition parameters as for CRISPR-RCA and Alt-R CRISPR-ATTO550 telomere imaging (Figure 6 in the main manuscript). The 'Merge' panel depicts a merged view of the nuclear DAPI staining (blue) and background fluorescence contribution by the CRISPR-RCA (i.e., $DNA_{RCA}$-ATTO550, top panel) or Alt-R CRISPR-ATTO550 (bottom panel).

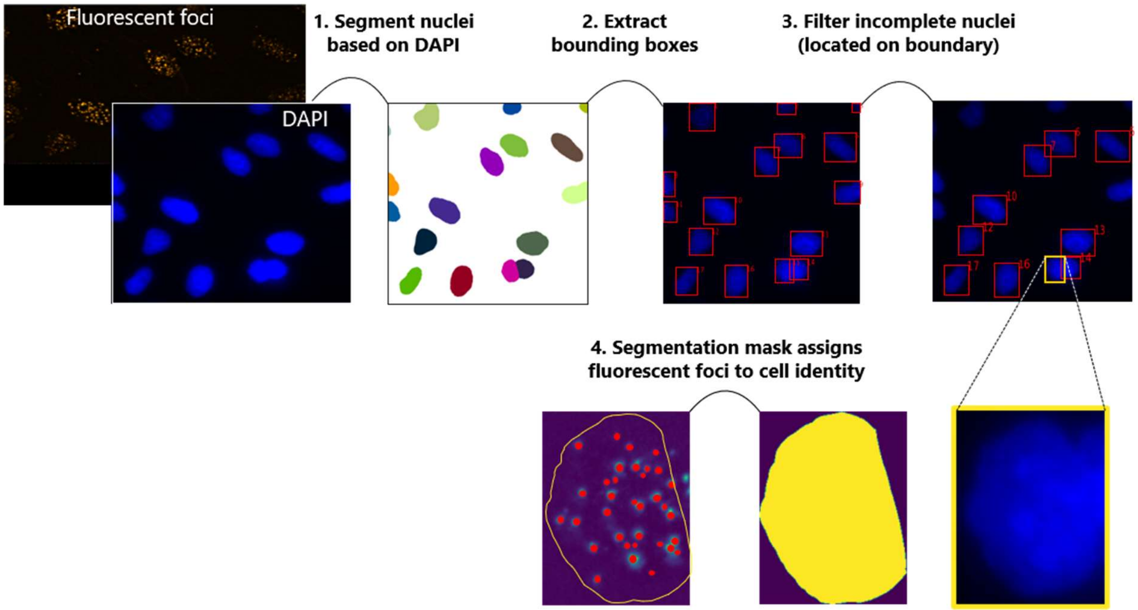


**Figure S10. Auxiliary schematic of image processing workflow.** See Materials and Methods 'Nucleus segmentation for fluorescent foci detection'.

**Table S1. Overview of nucleic acid sequences used in this research**. Protospacer sequences are underlined in the crRNAs and the dsDNA target and PAM sequence in bold. Alt-R modifications are IDT proprietary modifications. For cloning primers, bases containing the primer overhangs for ligation to the destination vector are in lowercase.

| **Name** | **Sequence (5'-3')** |
|---|---|
| gBlock dsDNA | TGGCGGCCTAAGGACTCGGAGTTCCGGAAGTGGCCAGTACGTGGACGA<br>CTTCGGCTCACAGTGAGTCTGAGGGTTAGGGTTAGGGTTAGGGTTA**GG**<br>**G**TGGATGATGATATCGCAACGCTCGTCGTCGACAACGGCTCCAGCATGT<br>GCAAGGCTAGCTTCGCGTTCGACGATGCACCTCGGGCCGTCTTCCCCTC<br>CATCGTGGGGCGCTTCAGGCACCAGGTATGGTAGCTGGCTGAGTGTGG<br>CAGTCACGTGAGCGATCGGGAAGCAAGTTCGCTTTCTCTGCACAGGAG<br>CCTCCCGGTTTCCGGAGTGTTGGCTGCGCCCGTGCTCAGGGCTTCTTGT<br>CCTATCCTTCCCAGGGCGTGATGGTGGGCATGGGTCAGAAGATTCCTAT<br>GTGGGCGACGAGGCCCAGAGCAAGAGAGGCATCCTCACCCTGAAG |
| Forward primer | TGGCGGCCTAAGGACTCG |
| Reverse primer | CTTCAGGGTGAGGATGCCTCTC |
| crRNA$_{Telomere}$ | GUUAGGGUUAGGGUUAGGGUUAGUUUUAGAGCUAUGCUGUUUUG |

| | |
|---|---|
| crRNA$_{GAL4}$ | GAACGACUAGUUAGGCGUGUAGUUUUAGAGCUAUGCUGUUUUG |
| Alt-R tracrRNA-ATTO550 | AGCAUAGCAAGUUAAAAUAAGGCUAGUCCGUUAUCAACUU GAAAAAGUGGCACCGAGUCGGUGCUUU |
| tracrRNA$^{5'Azide}$ | /5Azide/AGCAUAGCAAGUUAAAAUAAGGCUAGUCCGUUAU CAACUUGAAAAAGUGGCACCGAGUCGGUGCUUU |
| DBCO-TEG-DNA-ATTO550 | /5DBCOTEG/TGGTGAGGCTAGCTACAACGAGGTTGT |
| DNA$_{Hyb}$-ATTO488 | ACAACCTCGTTGTAGCTAGCCTCACCA/3ATTO488N/ |
| DBCO-TEG-RCA primer | /5DBCOTEG/TGCGAAGCGTGGATTCATTG |
| Dumbbell RCA template | GTTACCTTTCAATGAATCCACGCTTCGCACGGTTGGTGTAACGCTTGGA CTTTTCGGATTTCTAGGATCTCCAAGTATGTTCTAAGTCCAAGC |
| DNA$_{RCA}$-ATTO550 | GGATTTCTAGGATCTC/3ATTO550N/ |
| Primer AA_CS_021 | tgtctcagctgggaggcgacggatcaATGGACAAAGATTGCG |
| Primer AA_CS_022 | tttgtggcggcaggtcgctttgatcctCCCAGACCCGGTT |
| Primer AA_CS_035 | Ataaggatccgagctcgagcatggacaagaagtacagcatcg |
| Primer AA_CS_036 | agccaagcttcgaattcttatCCCAGACCCGGTTTACCCA |
| RCA primer (in vitro RCA, non-clickable) | TGCGAAGCGTGGATTCATTG |
| BB$_{PrimerBinding}$ | GTTACCTTTCAATGAATCCACGCTTCGCACGGTTGGTGTAACGCTTG |
| BB$_{Readout}$ | GACTTTTCGGATTTCTAGGATCTCCAAGTATGTTCTAAGTCCAAGC |
| BB$_{Nonsense}$ | GACTTTTCTGAAGGTTCTGACGCACAAGTATGTTCTAAGTCCAAGC |
| RCA template$_{Nonsense}$ | GTTACCTTTCAATGAATCCACGCTTCGCACGGTTGGTGTAACGCTTGGA CTTTTCTGAAGGTTCTGACGCACAAGTATGTTCTAAGTCCAAGC |

**Table S2. Overview of fluorescent foci detection parameters utilized per analyzed dataset.**

| Dataset (Corresp. figure) | Filter | Gauss | mn1 | mx1 | thresh1 | mn2 | mx2 | thresh2 |
|---|---|---|---|---|---|---|---|---|
| Figure 3A, tracrRNA-click | 0 | 1 | 2 | 3 | 0.004 | 0 | 0 | 0 |
| Figure 3A, gRNA-click | 0 | 1 | 2 | 3 | 0.004 | 0 | 0 | 0 |
| Figure 3A, RNP-click | 0 | 1 | 2 | 3 | 0.004 | 0 | 0 | 0 |
| Figure 3D, CRISPR | 0 | 1 | 2 | 3 | 0.002 | 2 | 3 | 0.002 |
| Figure 3D, PNA-FISH | 0 | 1 | 2 | 3 | 0.002 | 2 | 3 | 0.002 |
| Figure 3E, tracrRNA-click | 0 | 0 | 2 | 4 | 0.004 | 2 | 4 | 0.004 |
| Figure 3E, no tracrRNA-click | 0 | 0 | 2 | 4 | 0.004 | 2 | 4 | 0.004 |
| Figure 5A, tracrRNA-click | 0 | 1 | 2 | 3 | 0.005 | 0 | 0 | 0 |
| Figure 5A, gRNA-click | 0 | 1 | 2 | 3 | 0.006 | 0 | 0 | 0 |
| Figure 5A, RNP-click | 0 | 1 | 2 | 3 | 0.002 | 0 | 0 | 0 |
